# The viral SUMO-targeted Ubiquitin Ligase ICP0 is phosphorylated and activated by host kinase Chk2

**DOI:** 10.1101/709485

**Authors:** Dambarudhar SS Hembram, Hitendra Negi, Poulomi Biswas, Vasvi Tripathi, Lokesh Bhushan, Divya Shet, Vikas Kumar, Ranabir Das

**Affiliations:** National Center for Biological Sciences, TIFR, Bangalore, India; SASTRA University, Thirumalaisamudram, Thanjavur, India; Indian Institute of Technology, Kharagpur; Raman Research Institute, Bangalore; University of Nebraska Medical Center, Omaha, NE

**Keywords:** PML Nuclear Bodies, Transcriptional regulation, Ubiquitination, SUMOylation, Phosphorylation, Host-Virus Interactions, NMR spectroscopy

## Abstract

When the Herpes Simplex virus (HSV) genome enters the nucleus for replication and transcription, phase-segregated nuclear protein bodies called PML Nuclear Bodies (PML NBs) colocalize with the genome and repress it. HSV encodes a SUMO-targeted Ubiquitin ligase ICP0 that degrades PML NBs to alleviate the repression. The molecular mechanism used by ICP0 to target PML NBs is unclear. For reasons unknown, the growth of HSV is dependent on the ATM/Chk2 pathway. Here we identify a bonafide SUMO-Interacting motif in ICP0 (SLS4) that is essential and sufficient to target SUMOylated proteins in PML NBs like PML and Sp100. Phosphorylation of SLS4 creates new salt-bridges between SUMO and SLS4, increases the SUMO/SLS4 affinity and switches ICP0 into a potent STUbL. We also report that ICP0 exploits the kinase Chk2 to phosphorylate SLS4 and enhance its STUbL activity. Our results uncover how a viral STUbL counters antiviral response by exploiting an unprecedented mechanism involving three post-translational modifications; ubiquitination, SUMOylation, and phosphorylation.

## Introduction

DNA viruses optimize the host nuclear environment for genome replication and transcription. PML nuclear bodies (PML NBs, also known as ND10 or PODs) are dynamic nuclear substructures located in the interchromosomal space. These multiprotein phase-segregated bodies serve as an antiviral response to prevent transcription of viral genes during infection (Ascoli and Maul, 1991; Stuurman *et al*., 1992; Lallemand-Breitenbach and de Thé, 2010; Banani *et al*., 2016). In the nucleus, incoming viral genome triggers the spontaneous formation of PML NBs around it, which then transcriptionally represses the genome (Everett *et al*., 2004, 2006, 2007)(Tavalai and Stamminger, 2009). PML NBs inhibit the growth of several viruses, including the entire *herpesviridae* family, adenovirus, papovavirus SV40, papovavirus AAV, and polyomavirus (Everett, 2001). Viruses counteract by disrupting and degrading the PML NBs soon after their entry into the host nucleus (Tavalai and Stamminger, 2009). Herpes Simplex Virus-1 (HSV-1) expresses a ubiquitin ligase (E3) known as the Infected Cell Polypeptide 0 (ICP0) in the immediate early stages of the *lytic* cycle (Maul, Guldner and Spivack, 1993). A prominent role of ICP0 is to degrade PML NBs via the ubiquitin-proteasome pathway (Chelbi-alix and The, 1999; Parkinson and Everett, 2000). ICP0 is a RING-finger E3, and its catalytic RING-finger domain is essential for the degradation of PML NBs (Everett and Maul, 1994). If ICP0 is deleted, HSV-1 fails to degrade PML NBs, resulting in reduced viral gene expression (Maul, Guldner and Spivack, 1993; Everett *et al*., 2008). These results suggest that degradation of PML NB is imperative for HSV-1, and ICP0 is indispensable for this purpose.

Interestingly, another ubiquitin-like molecule, the Small Ubiquitin-like Modifier (SUMO) is also involved in the function of ICP0. The essential constituent proteins of PML NBs such as PML, hDaxx, and Sp100 are heavily SUMOylated (Lallemand-Breitenbach and de Thé, 2010). A possible mechanism for ICP0 to disrupt PML NBs would be to target these SUMOylated proteins. E3s that target SUMOylated substrates are known as SUMO-Targeted Ubiquitin Ligases (STUbLs)(Prudden *et al*., 2007). Apart from the catalytic RING-finger domain, STUbLs contain SUMO-Interacting-Motifs (SIMs), which bind to SUMOylated substrates and enable the E3 to assemble polyubiquitin chains on the substrates. Recent investigations indicate that indeed ICP0 is a viral STUbL, which recognizes the SUMOylated proteins by one or more of its seven predicted SIM-like-sequences (SLS1-7) (Boutell and Everett, 2013; Everett *et al*., 2014a). However, the identity of the bonafide SIMs in ICP0 and the molecular details of how ICP0 targets SUMOylated substrates remain elusive. Intriguingly, PML NBs are short-lived after ICP0 expression, indicating that they undergo rapid degradation due to ICP0. Typically SUMO/SIM interactions are weak, with dissociation constants around 100 μM (Song *et al*., 2004). It is unclear how ICP0 can efficiently target SUMOylated substrates for rapid ubiquitination despite the weak affinity of SUMO/SIM interaction.

The viral genome also activates other antiviral responses such as the DNA damage response (DDR) to cap the foreign genome. Viruses have evolved to either inhibit or circumvent such responses (Chaurushiya and Weitzman, 2009; Weitzman and Weller, 2011). Interestingly, in some cases, these responses benefit viral growth. For example, HSV, polyomavirus, papovavirus SV40 directly activate DDR, and their growth declines in the absence of DDR proteins (Everett, 2006; Chaurushiya and Weitzman, 2009). However, why these proteins benefit viral growth is unclear. During HSV-1 infection, ICP0 activates the ATM/Chk2 pathway, which is a key component of DDR. The activation probably occurs at the PML NBs, where ICP0, ATM, and Chk2 localize transiently (Dellaire *et al*., 2006; Li *et al*., 2008). Chk2 deletion severely reduced the growth of HSV-1, a phenotype similar to ICP0-deleted HSV-1, signifying the relevance of Chk2 for viral growth (Li *et al*., 2008). Given that ATM/Chk2 is a component of DDR (an antiviral response), the purpose behind its activation is rather non-intuitive.

We carried out a systematic investigation of interactions between SUMO1 and SUMO2 with all the seven predicted SLS regions (SLS1-SLS7) by NMR. The results demonstrate that SLS4 is the sole bonafide SIM in ICP0. Structures of the SUMO/SLS4 complexes studied here uncover a set of hydrophobic and electrostatic contacts between SUMO and SLS4, which are critical for targeting SUMOylated substrates like PML and Sp100 for ubiquitination. Interestingly, ICP0 ubiquitinates both the SUMO and the substrate in a SUMOylated substrate, a mechanism that could be crucial to prevent regulation by host deSUMOylating enzymes. Two phosphoserines adjacent to SLS4 is phosphorylated in cellular conditions. Phosphorylation of SLS4 enhances its affinity for SUMO by several folds, resulting in enhanced STUbL activity. Further structural and biochemical studies confirm that two kinases CK1 and Chk2 function in tandem to phosphorylate SLS4 and enhance the STUbL activity of ICP0. Altogether, our results demonstrate how a viral E3 exploits the cross-talk of three host post-translational modifications (PTMs), ubiquitination, SUMOylation, and phosphorylation to degrade host antiviral responses efficiently.

## Results

### SLS4 is the sole bonafide SIM in ICP0

One or more of the seven predicted SLS domains in ICP0 (SLS1-SLS7) could be responsible for targeting SUMOylated substrates (Figure 1A). Hence, all seven domains were tested for binding to SUMO. Seven peptides were designed corresponding to the SLS domains (Table S1). The affinity of the SIM domains for SUMO is typically weak (Song *et al*., 2004). Since NMR spectroscopy can detect binding over a broad range of affinities including very weak affinities, the binding of SLS domains to SUMO1 and SUMO2 was tested by NMR. SLS4 was titrated into a sample of ^15^N isotope-labeled SUMO1, and the effect on SUMO1 was detected by ^15^N-edited Heteronuclear Single Quantum Coherence (HSQC) experiments. Perturbations due to the altered chemical environment upon ligand binding induce changes in the chemical shift of the backbone amide resonances (Figure 1B). The chemical shift perturbations (CSP) plotted in Figure 1C shows that the maximal perturbations upon binding SLS4 occur in the region between β2 and α1 of SUMO1 (Figure 1D), which was identified as the binding site of SLS4. SIMs from PML, RNF4, and PIASX bind to a similar region in SUMO (Song *et al*., 2004; Xu *et al*., 2014; Tremblay-belzile *et al*., 2015). The same titration experiments were repeated for other SLS domains. However, none of the other SLS domains showed any consistent peak shifts in the SUMO1 spectra upon titration (an example titration of SLS7 is given in Figure S1A), indicating that the other SLS domains do not bind SUMO1. The shifts in amide resonances in the SUMO1/SLS4 titration spectra was fit against the ligand:protein concentration ratio to yield the K_d_ of 128(±13) μM (Figure S2A).

**Figure 1.**
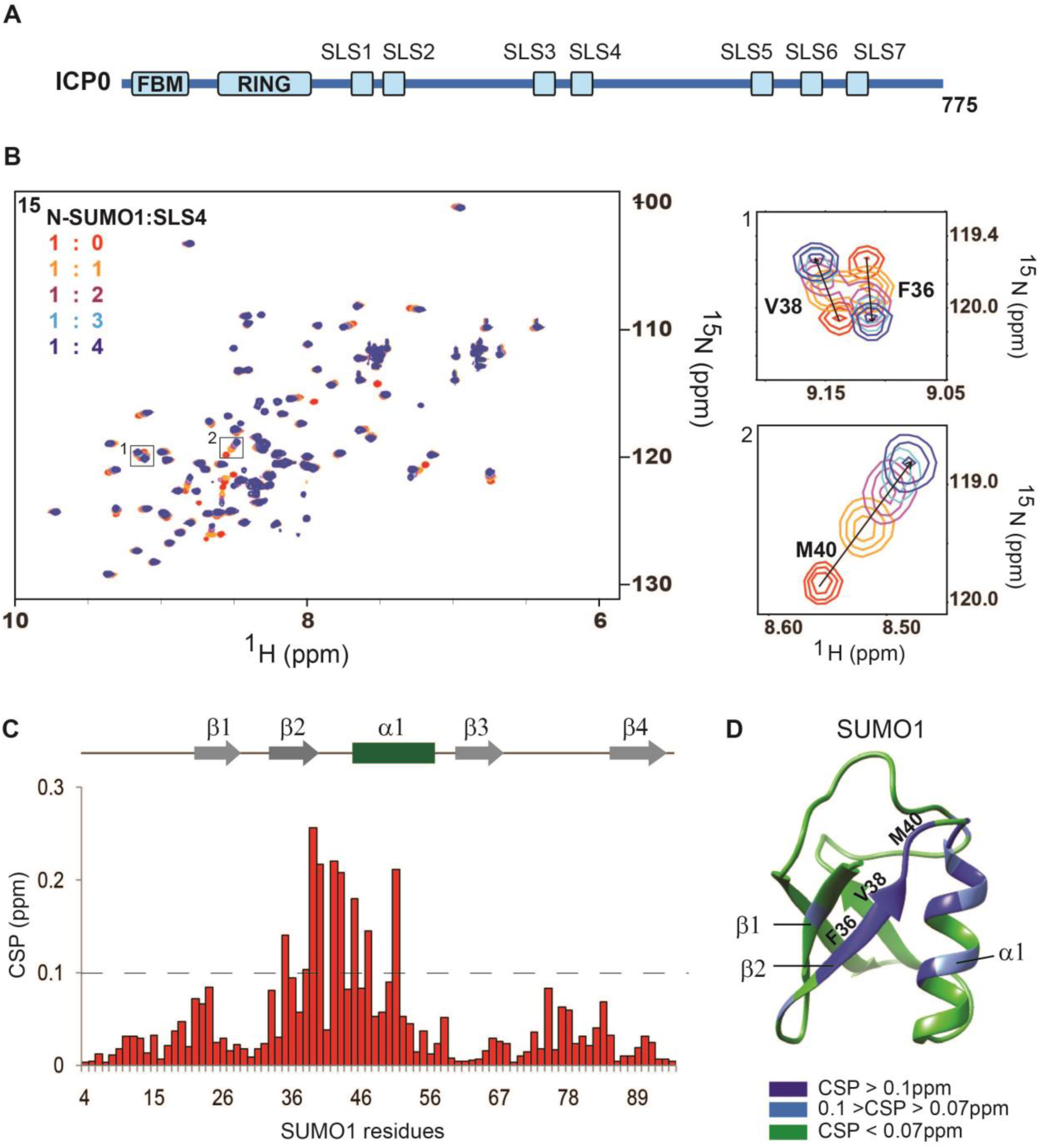
Interactions between SLS4 and SUMO1 studied by NMR. (A) The domains in ICP0 relevant to this study. The FHA Binding Motif (FBM), the RING finger domain, and the SLS domains are shown. (B) Overlay of the ^15^N-edited HSQC spectra of free SUMO1 (red) with different stoichiometric ratios of SLS4 as given in the top left-hand side of the spectra. Two regions of the spectra are expanded to show SUMO1 peaks shift upon titration with SLS4. The chemical shift perturbations (CSP) between the free and the bound form are calculated as CSP = [(δ^H^_free_ – δ^H^_bound_)^2^+ ((δ^N^_free_ – δ^N^_bound_)/5)^2^]^1/2^, where δ^H^ and δ^N^ are the chemical shift of the amide hydrogen and nitrogen, respectively. (C) The CSPs for each residue in SUMO1 upon binding to SLS4. The black dashed line indicates 2*standard deviation. The residues with CSPs significantly above the dashed line are probably at the interface of the SUMO1/SLS4 complex. The secondary structure alignment of SUMO1 against its sequence is provided above the plot. (D) Significant CSPs mapped on the SUMO1 structure. The residues corresponding to the expanded spectra in (B) are labeled on the structure.

The titration experiments were repeated with ^15^N-labeled SUMO2, SLS4, and monitored using the ^15^N-edited HSQC spectra of SUMO2 (Figure 2A). Apart from a few long-range CSPs in β1 beta-strand of SUMO2, the majority of CSPs occurred in the region between β2 and α1(Figure 2B and 2C). None of the other SLS domains induced consistent shifts in the SUMO2 amide resonances upon titration (Figure S1B). The shifts in amide resonances in the SUMO2/SLS4 titration spectra was fit against the ligand:protein concentration ratio to yield the K_d_ of 81(±17) μM (Figure S2B). It has been indicated recently that the region, including the SLS domains SLS5, SLS6, and SLS7, could be important for binding SUMO (Everett *et al*., 2014a). Hence, SUMO1 and SUMO2 were titrated against a peptide that included SLS5, 6 and 7 (SLS567). However, SLS567 did not bind SUMO2 (Figure S3A and S3B) and interacted very weakly to SUMO1 (Figure S3C). Since SLS4 is the only SLS domain in ICP0 that binds SUMO1 or SUMO2, it was identified as the sole bonafide SIM in ICP0.

**Figure 2.**
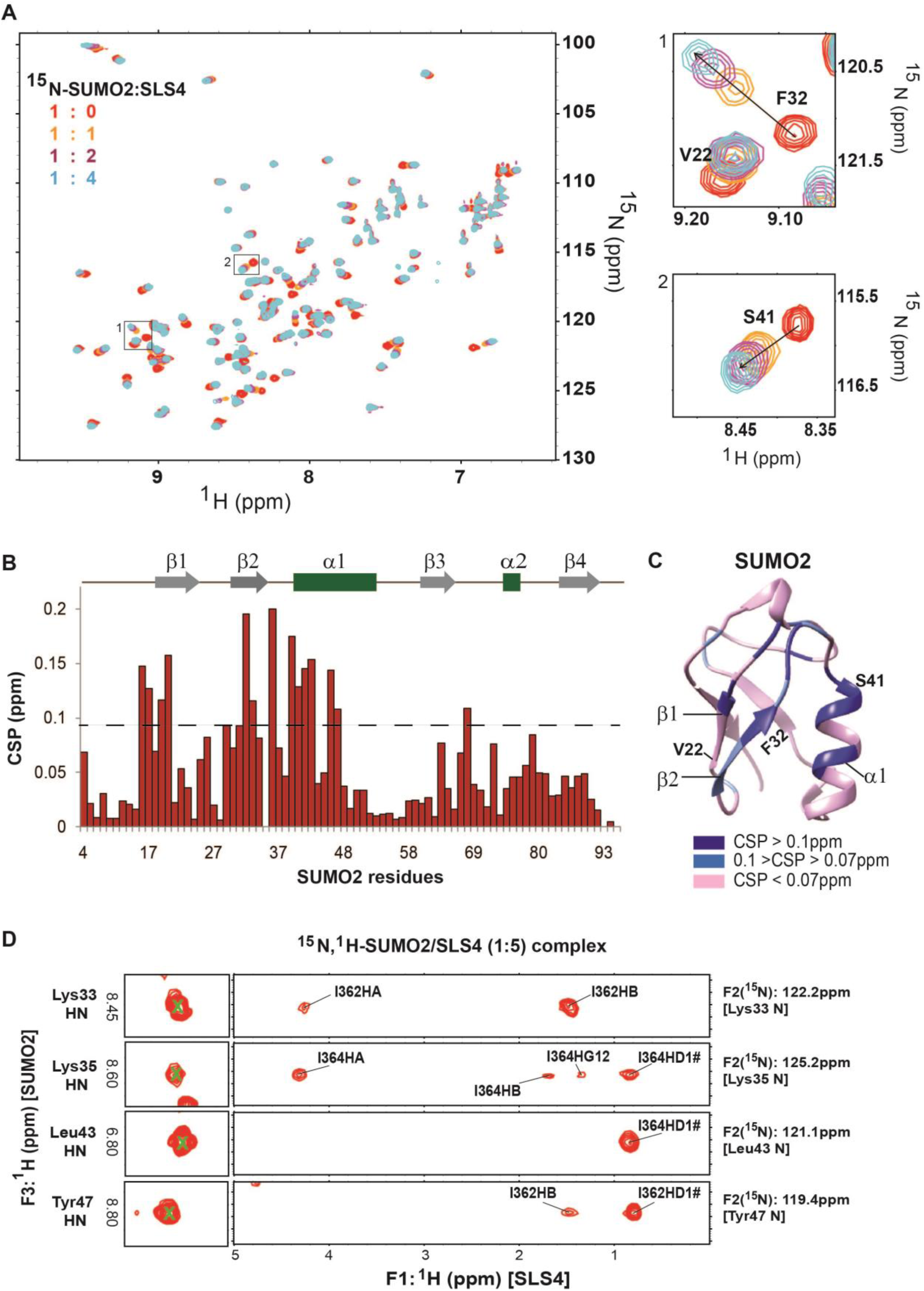
Interactions between SLS4 and SUMO2 studied by NMR. (A) Overlay of the ^15^N-edited HSQC spectra of free SUMO2 (red) with different stoichiometric ratios of SLS4 as given in the top left-hand side of the spectra. Two regions of the spectra are expanded to show SLS4 induced shifts in the SUMO2. The chemical shift perturbations (CSP) between the free and the bound form are calculated as given in Figure 1 legend. (B) The CSPs observed in SUMO2 upon binding SLS4 are shown. The dashed line shows the 2*standard deviation. (C) Significant CSPs mapped on the SUMO2 structure. The residues corresponding to the expanded spectra in (A) is labeled on the structure. (D) Selected strips from the ^15^N-edited NOESY-HSQC spectra depicting intermolecular NOEs between amide protons of SUMO2 and protons of the SLS4 domain. ^15^N and ^1^H assignment of SUMO2 amide atoms are given on the right and left of the strips, respectively. The protons of SLS4 that show NOEs to SLS4 are assigned.

### Structure of SUMO1/SLS4 and SUMO2/SLS4 complexes

The structure of SUMO1/SLS4 and SUMO2/SLS4 complexes were determined to obtain molecular details of their interaction. 2D ^1^H-^1^H TOCSY and ^1^H-^1^H NOESY experiments provided the proton chemical shifts of SLS4. Although the SLS4 peptide used in the experiments was twenty amino acids long, only the central twelve amino acids (N357-P368) exhibited resonances in the amide region in the ^1^H-^1^H TOCSY and ^1^H-^1^H NOESY spectra of the peptide, indicating that the N- and C-terminal regions of SLS4 are dynamic, and their resonances are broadened due to conformational exchange. ^15^N-edited NOESY-HSQC experiment was carried out on a ^15^N, ^2^H-labeled-SUMO2/SLS4 (1:5) complex to obtain intermolecular NOEs between SUMO2 and SLS4 (Figure 2D). The structure of SLS4 was determined using torsion angles from ^1^Hα assignments and ^1^H-^1^H NOESY restraints. Using the SLS4 structure, SUMO2 structure (PDB: 1WM3), and the intermolecular NOEs as distance restraints; the structure of the SUMO2/SLS4 complex was determined by HADDOCK (Dominguez, Boelens and Bonvin, 2003). The refinement statistics are provided in Table 1. Figure S4A shows the twenty lowest energy structures, which superimposed with a root-mean-square deviation (rmsd) of 0.6 Å. The lowest energy structure given in Figure 3A verified that SLS4 binds to the region between β2 and α1 as predicted by CSPs from the titration experiment. SLS4 forms a β-strand parallel to the β2 strands of SUMO2.

**Figure 3.**
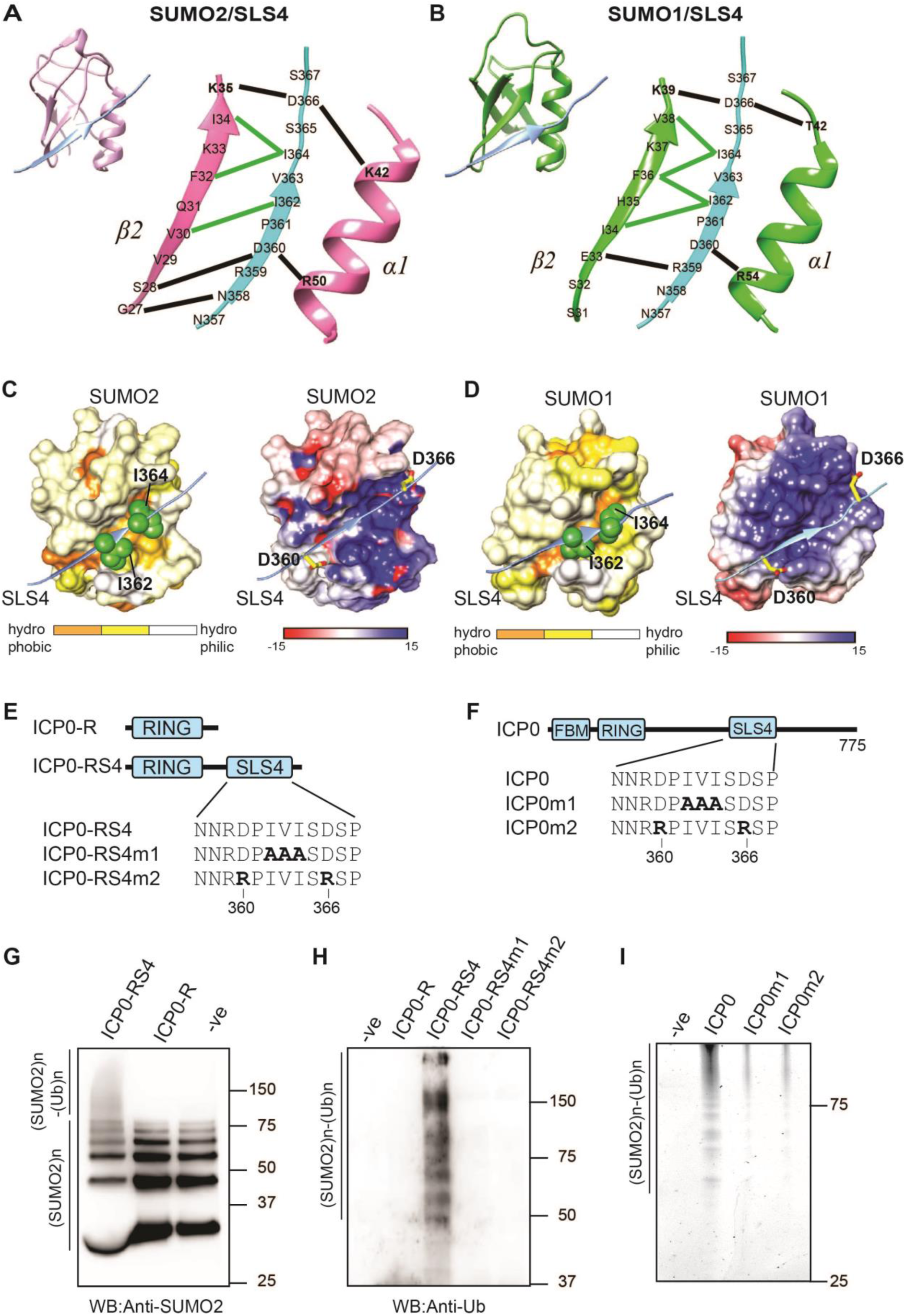
The hydrophobic and electrostatic contacts at the SUMO/SLS4 is essential for the STUbL activity of ICP0. (A) and (B) Left-hand corner represents the lowest energy structure of the SUMO2/SLS4 and SUMO1/SLS4 complex, respectively. SUMO2 is colored in pink, SUMO1 is colored in green, and SLS4 is colored in light blue. The interface between SLS4 and SUMO1/2 is expanded on the right. The salt-bridges between SLS4 and SUMO1/2 are shown as black lines. The hydrophobic contacts between the central PIVI residues in SLS4 and SUMO1/2 are shown as green lines. (C) Surface representation of hydrophobic regions of SUMO2 in the SUMO2/SLS4 complex is shown in the left side. The surfaces are colored according to hydrophobicity scale given below. The side-chains of Ile362 and Ile364 in SLS4 are shown in green. The electrostatic surface of SUMO2 in the SUMO2/SLS4 complex is shown on the right side. Positive to negative charges are shown by blue to red. The ABSF program calculated the electrostatic surface potential. The acidic residues of SLS4, Asp360, and Asp366 are shown in yellow. The side-chain oxygen atoms are colored in red. (D) Same as (C) shown for SUMO1 in the SUMO1/SLS4 complex. (E) ICP0-R, ICP0-RS4, and its mutants used in this study. (F) ICP0 and its mutants used in this study. (G) Ubiquitination assays using Ube1 as E1, Ube2d1 as E2, and either ICP0-R or ICP0-RS4 as the E3, SUMO2-chains as the substrate and blotted with the anti-SUMO2 antibody. (H) Similar ubiquitination reaction as in (G) carried out on SUMO2-affinity beads, washed and blotted with the anti-Ub antibody. (I) Similar ubiquitination reaction as in (H) using full-length ICP0 and its mutants as the E3, and, Alexa-Ub. The (-ve) control lane in (G), (H) and (I) lacks the E3.

**Table 1.**
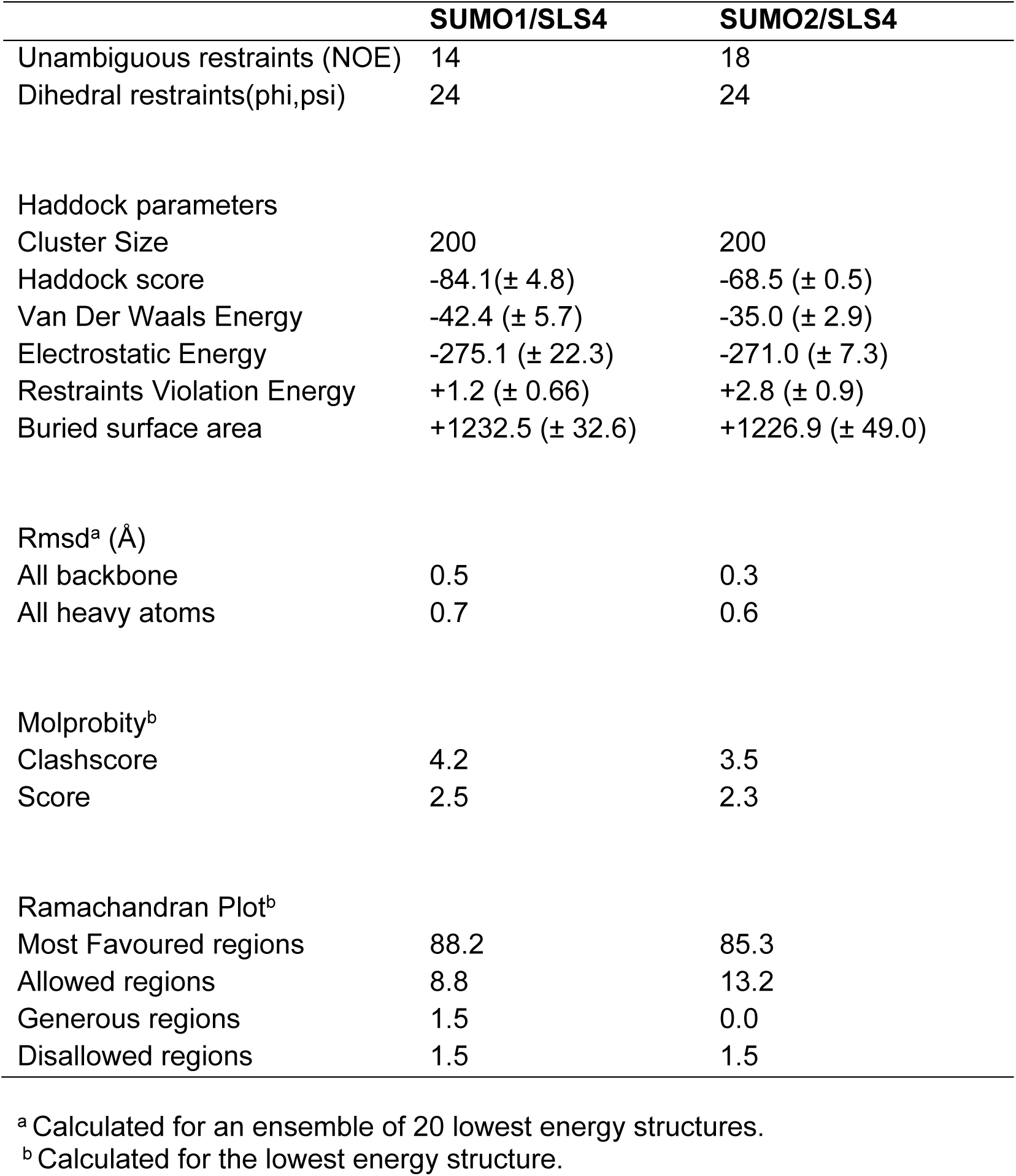
NMR and refinement statistics of the SUMO/SLS4 complexes.

Both the hydrophobic and electrostatic interactions contribute to the binding between SUMO2 and SLS4. A set of hydrophobic contacts are observed between the central SLS4 residues (PIVI) and the β2-strand of SUMO2 (Figure 3A, Figure S4B, Table S2). A hydrophobic shallow groove is present between β2 and α1 in SUMO2, and the hydrophobic side-chains of the SLS4 reciprocate this groove (Figure 3C). SUMO2 has a significant positively charged surface at the SIM binding interface (Figure 3C), which complements the negatively charged residues in SLS4. Hence, a network of salt-bridges form at the N- and C-termini of central SLS4 region, which stabilizes the SUMO2/SLS4 interaction (Figure 3A).

^15^N-edited NOESY-HSQC on a sample of ^15^N, ^2^H-SUMO1/SLS4 (1:5) complex detected the intermolecular NOEs between SUMO1 and SLS4 (Figure S5A). The structure of the SUMO1/SLS4 complex was determined using the measured intermolecular NOEs (Table 1). The twenty lowest energy structures superposed with an rmsd of 0.7 Å (Figure S5B). The lowest energy structure given in Figure 3B shows that SLS4 binds to the region between β2 and α1 in SUMO1 as predicted by the CSPs. The packing of hydrophobic surfaces and the surface charge complementation were also evident in the SUMO1/SLS4 complex (Figure 3D and S5C, Table S3).

### The SLS4 and the RING finger domain is sufficient for STUbL activity of ICP0

STUbLs include two essential domains, the RING finger domain that interacts with the E2∼Ub conjugate, and SIM domains that interact with the SUMOylated substrates (Prudden *et al*., 2007; Sriramachandran and Dohmen, 2014). ICP0 includes both the catalytic RING finger and the bonafide SIM SLS4. Two chimeras were prepared to investigate if these two domains in ICP0 are sufficient for its STUbL activity; ICP0-R and ICP0-RS4. ICP0-R includes the RING finger domain of ICP0, and, ICP0-RS4 includes both the RING finger domain and the SLS4 domain (Figure 3E). A ubiquitination reaction with Ube1 (E1), Ube2d1 (E2), Ub, and either ICP0-R or ICP0-RS4 were carried out at 37°C. Both the chimeras could catalyze polyubiquitin chains owing to the presence of the RING finger domain (Figure S6A). Then the STUbL activity of these chimeras was examined by their ability to ubiquitinate SUMO2-chains (as a substitute of the SUMOylated substrate). The ubiquitination reaction was repeated in the presence of SUMO2 chains. Whereas ICP0-RS4 could identify SUMO2-chains and assemble polyubiquitin chains on them, ICP0-R could not (Figure 3G and 3H). When the central hydrophobic Ile362-Val363-Ile364 residues in SLS4 were mutated to alanines, the ubiquitination activity of ICP0-RS4 was unaffected, but its STUbL activity diminished (Figure 3H and Figure S6B). A similar effect was observed when the negatively charged residues flanking the central region Asp360 and Asp366 were mutated to positive charges (Figure 3H). When the experiments were repeated with the full-length ICP0 and its mutants (Figure 3F and Figure S6D), the results were similar (Figure 3I), indicating that mechanism of STUbL activity is similar between the full-length ICP0 and the ICP0-RS4 chimera. Altogether, these results demonstrate that the RING domain and SLS4 are necessary and sufficient for the STUbL activity of ICP0. Moreover, both the electrostatic and hydrophobic interactions between SUMO and SLS4 are important for STUbL activity.

### ICP0 ubiquitinates both the substrate and SUMO in the poly-SUMOylated substrate

Although ICP0 can identify and ubiquitinate free (SUMO)_n_ chains in solution, further experiments were carried out to assess if it can identify and ubiquitinate SUMOylated substrates. Three substrates were used to test the function of ICP0 (Figure 4A). The first substrate was a short region of PML (PMLs, aa:485-495), which includes the consensus SUMOylation site in PML K490 (Brand, Lenser and Hemmerich, 2010). The second substrate was a short region of PML NB constituent protein Sp100 (Sp100s, aa:292-305), which includes its consensus SUMOylation site K297 (Seeler *et al*., 2001). The third substrate is a long stretch of amino acids in the N-terminal region of PML (PML, aa:370-570), which includes the consensus SUMOylation site K490 and multiple Ubiquitination sites as predicted by UbPred and UbiSite (Tung and Ho, 2008; Akimov *et al*., 2018). A SUMOylation reaction with either FLAG-PMLs or FLAG-Sp100s confirmed that they are indeed SUMOylated *in-vitro* (Figure 4B). The SUMOylation reaction was captured on anti-FLAG affinity beads and washed to get rid of SUMOylation enzymes and free SUMO2 (Figure S6E). Then a ubiquitination reaction was carried out on the beads using Ube1, Ube2d1, ICP0, and Alexa-Ub. Following the reaction, EDTA was used to quench the reaction; the beads were washed thoroughly to get rid of the ubiquitination enzymes and any free polyubiquitin chains in solution. Following the wash, the beads containing ubiquitinated-SUMOylated substrate ((Ub)_n_∼(SUMO)_n_∼FLAG-PMLs) was detected. While ICP0 could identify and ubiquitinate SUMOylated FLAG-PMLs, the mutants ICP0m1 and ICP0m2 could not, validating that the electrostatic and hydrophobic interactions at the SUMO/SLS4 interface are critical for the STUbL activity (Figure 4C). Similarly, SUMOylated FLAG-Sp100s was ubiquitinated by ICP0, but not by ICP0m1 or ICP0m2 (Figure 4D).

**Figure 4.**
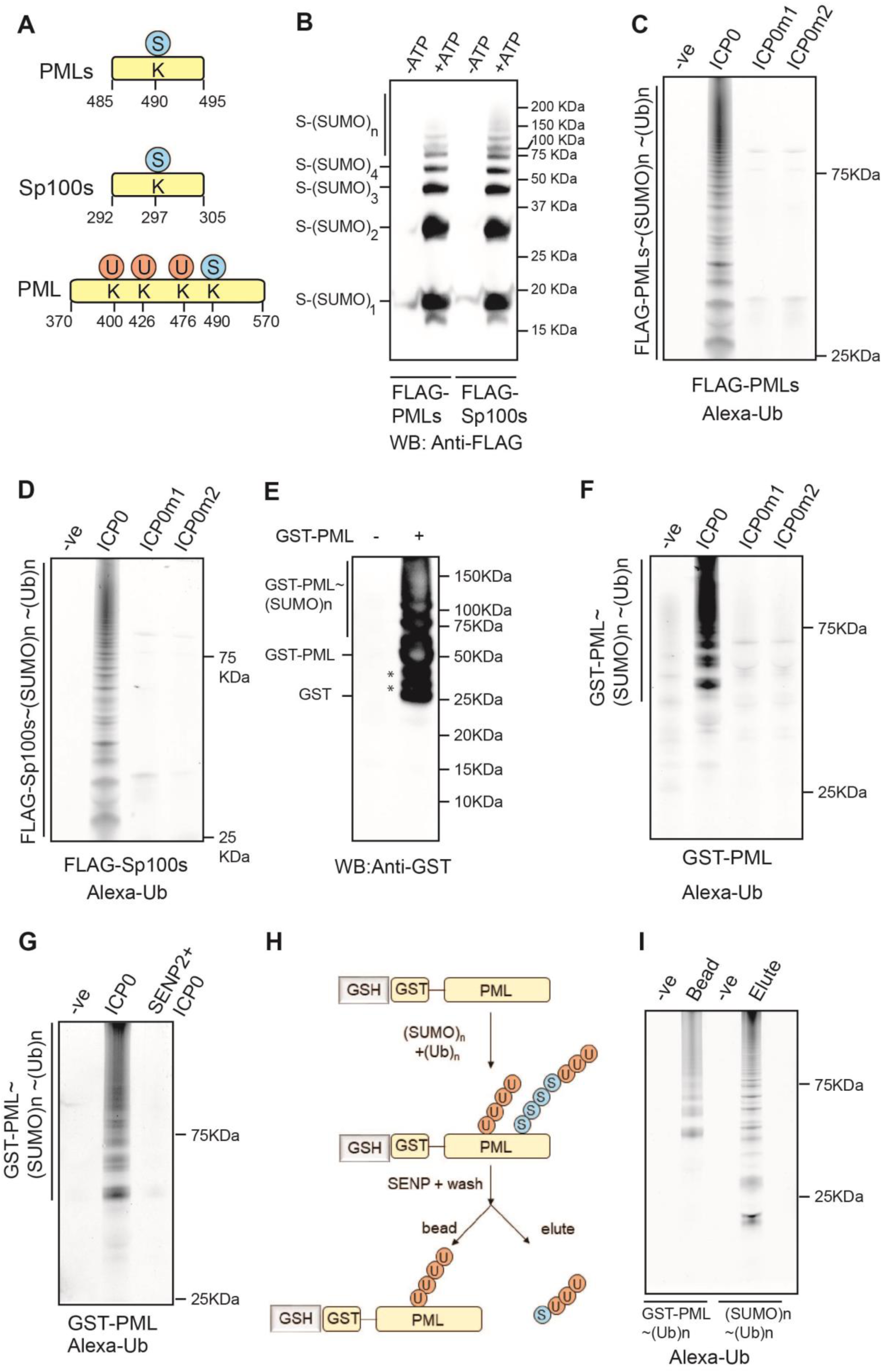
Ubiquitination of SUMOylated substrates by ICP0. (A) Schematic of the substrates PMLs (aa: 485-490 of protein PML), Sp100s (aa: 292-305 of protein Sp100) and PML (aa: 370-570 of protein PML). PMLs and Sp100s include the SUMOylation sites, while PML has both the SUMOylation site and predicted Ubiquitination sites in PML protein. (B) SUMOylation of PMLs and Sp100s. SUMOylation reactions were carried out with FLAG-tagged PMLs and Sp100s as the substrates, separated on SDS gel and blotted with Anti-FLAG antibody. (C) SUMOylated PMLs was captured on anti-FLAG affinity beads, washed (3x), and used as substrates in a ubiquitination reaction using Ube2d1 as the E2, Alexa-Ub, and either ICP0 or its mutants as the E3. Following the reaction, beads were washed (3x), separated on SDS gel and visualized in Uvitec (Cambridge). (D) Same as (C) using SUMOylated Sp100s. (E) SUMOylation reaction using GST-PML as the substrate. (F) Same as in (C), except the SUMOylated PML was captured on glutathione (GSH) beads. (G) Same as in (F), except that in one reaction (rightmost lane), SENP2 was added for two hours to cleave the SUMO chains on PML before adding ICP0 and other components of ubiquitination reaction. (H) Schematic of the experiments used to detect if the poly-Ub chains are assembled on SUMO chains or the substrate PML. (I) The beads and elutes of (H) are separated on SDS gel and visualized. The (-ve) lanes in (C), (D), (F), (G) and (I) do not include the SUMO activating enzyme SAE1/2.

Much like the shorter region of PML, the longer region of PML fused to an N-terminal GST tag was SUMOylated *in-vitro* (Figure 4E). The SUMOylation was not on the N-terminal GST (Figure S6F). The SUMOylation reaction was captured on Glutathione agarose beads and washed to get rid of SUMOylation enzymes and free SUMO. Consequently, ubiquitination reaction was carried out using Ube1, Ube2d1, ICP0, and Alexa-Ub, washed, and the beads were separated on SDS gel for visualization. ICP0m1 and ICP0m2 could not ubiquitinate SUMOylated PML as efficiently as ICP0 (Figure 4F). If the SUMOylated PML was cleaved by deSUMOylating enzyme SENP2 and washed prior to ubiquitination, ICP0 could not ubiquitinate PML effectively, indicating that ICP0 specifically targets SUMOylated PML (Figure 4G). Another experiment was carried out to detect if ICP0 assembles polyubiquitin chains directly on the substrate PML or on the poly-SUMO chains conjugated to PML (Figure 4H). Post the ubiquitination reaction; the beads were washed, treated with SENP2, then spun to separate the GST-PML and SUMO chains, followed by detection. ICP0 ubiquitinates both the PML and SUMO (Figure 4I), suggesting a mechanism where the viral STUbL ensures that the substrate is marked for degradation and cannot be rescued by host deSUMOylating enzymes.

### Phosphorylated SLS4 binds SUMO1 and SUMO2 with higher affinity

Phosphorylation of ICP0 can significantly regulate its activity (Davido *et al*., 2005; Boutell *et al*., 2008; Mostafa *et al*., 2011). Interestingly, a variant of ICP0 that cannot be phosphorylated at Ser365, Ser367, and Ser371 lack the potency to degrade PML NBs (Boutell *et al*., 2008). When the phosphorylation of ICP0 was checked in HEK293T cells, Ser365, Ser367, and Ser371 were indeed phosphorylated (Figure 5A). In the SUMO/SLS4 structures, Ser371 is dynamic and far from the SUMO/SLS4 interface. On the other hand, both Ser365 and Ser367 are present at the SUMO/SLS4 interface. The effect of phosphorylation at 365 on SUMO/SLS4 interaction was tested by using a pSLS4 peptide, which is similar to SLS4 but phosphorylated at Ser365 (Figure 5B). These NMR titrations were carried out in conditions similar to the SLS4 titration. Figure S7A shows the pattern of CSPs observed in SUMO2, which is similar to CSPs observed during the titration with SLS4, implying that the interface of pSLS4 is similar to SLS4. However, the magnitude of average CSP was higher, indicating that a higher population of the SUMO2/pSLS4 was in bound conformation compared to SUMO2/SLS4 at the same protein to ligand ratio and implying that pSLS4 has a higher affinity for SUMO2 than SLS4. Indeed, upon fitting the titration data, the K_d_ was found to be 7(±2) μM compared to the 81(±17) μM observed for SLS4 (Figure 5C, Figure S7B).

**Figure 5.**
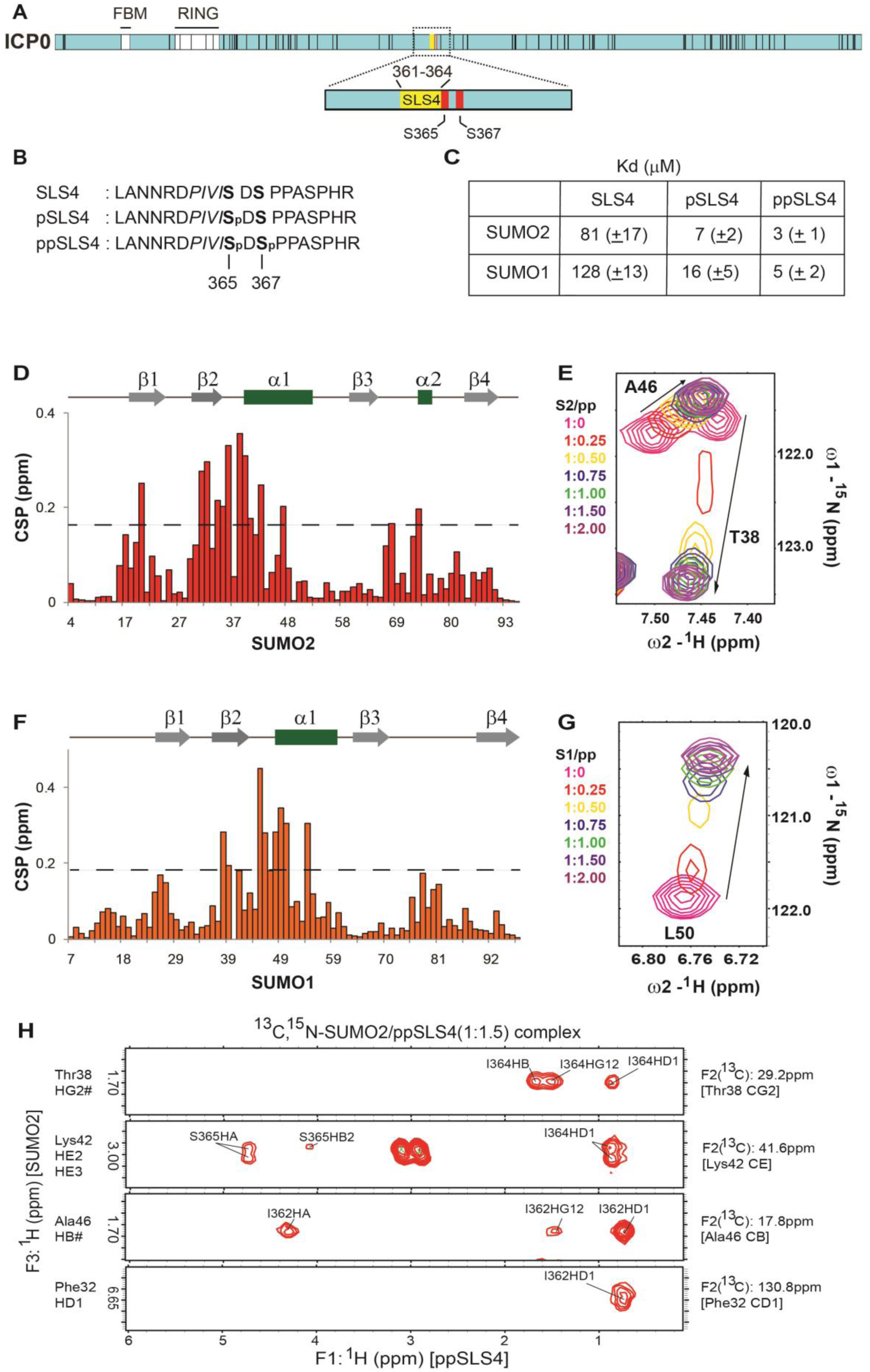
ICP0 is phosphorylated at SLS4, which increases its affinity for SUMO. (A) Phosphorylation of ICP0 detected in HEK293T by Mass-spectrometry. All vertical black lines denote detected phosphorylation sites in ICP0. The region around SLS4 is expanded to show that only two serines immediately adjacent to SIM1 are phosphorylated. Ser365 and Ser367 are colored as red lines. (B) The sequence of the SLS4, pSLS4, and ppSLS4. (C) Dissociation constants of the binding of SUMO with SLS4, pSLS4, and ppSLS4. ppSLS4 induced CSPs observed in SUMO2 and SUMO1 are plotted in (D) and (F), respectively. (E) and (G) shows a typical amide resonance that undergoes intermediate exchange when SUMO2 (S2/pp) and SUMO1 (S1/pp) bind ppSLS4, respectively. (H) Selected strips from the ^13^C,^15^N-filtered (F1), ^13^C,^15^N-edited (F2) NOESY HSQC spectra are depicting intermolecular NOEs between ^13^C-bonded protons of ^13^C, ^15^N-labeled SUMO2, and unlabeled ppSLS4. ^13^C and ^1^H assignment of SUMO2 atoms are given on the right and left of the strips, respectively. The protons of SLS4 that show NOEs to SUMO2 are labeled.

The titration was repeated with ppSLS4, where both Ser365 and Ser367 are phosphorylated (Figure 5B). The pattern of CSPs was similar, except for a slight increase in the average magnitude compared to pSLS4 (Figure 5D). A few peaks showed the pattern of intermediate exchange due to the increase in affinity (Figure 5E). The K_d_ was determined by fitting peak-shifts for the resonances that have less CSP and are still in fast-exchange. The K_d_ was determined to be 3(±1) μM (Figure 5C, Figure S9A).

When pSLS4 was titrated to SUMO1, the interface of pSLS4 was similar to SLS4 (Figure S8A). However, the average CSP was higher when compared to SLS4. Fitting the titration data yielded the K_d_ of 16(±5) μM (Figure S8B). When titrated with the diphosphorylated ppSLS4, the CSP pattern was conserved, but few peaks went into the intermediate exchange (Figure 5F and 5G). The peak shifts against SLS4 concentration were fit to yield the K_d_ of 5(±2) μM (Figure 5C, Figure S9B). Altogether, phosphorylation increased the affinity of SLS4 for SUMO by ∼25-fold.

### The phosphorylated side-chains of SLS4 make new electrostatic contacts with SUMO

The structure of the diphosphorylated SLS4 (ppSLS4) in complex with SUMO2 and SUMO1 was determined to understand the mechanism of the phosphorylation induced increase in SUMO/SLS4 affinity. A ^13^C,^15^N-filtered (F1), ^13^C,^15^N-edited (F2) NOESY HSQC was acquired on a sample of ^13^C, ^15^N-SUMO2/ppSLS4 at the stoichiometric ratio of 1:1.5 (SUMO2:ppSLS4) (Figure 5H). ^1^H-^1^H TOCSY and ^1^H-^1^H NOESY experiments provided the proton chemical shifts of ppSLS4. The chemical shifts of SUMO2 were assigned by comparing the ^15^N-^1^H edited HSQC and ^13^C-^1^H edited HSQC of ^13^C, ^15^N-SUMO2/ppSLS4 complex with the assignments of apo SUMO2. Using the intermolecular NOEs as distance restraints; the structure of the SUMO2/ppSLS4 complex was solved by HADDOCK (Dominguez, Boelens and Bonvin, 2003) (Table 2 provides the structural statistics). Figure S10A shows the twenty lowest energy structures, which superpose well with an rmsd of 0.7 Å. The structure of SUMO2/ppSLS4 was overall similar to SUMO2/SLS4. Two new hydrogen bonds formed the phosphorylated SLS4 side-chains and SUMO2 (Figure 6A). Lysine 42 of α1 in SUMO2 now forms hydrogen bonds with the phosphate oxygen atoms of pSer365 and pSer367. These hydrogen bonds contribute to the increased affinity between SUMO2 and ppSLS4.

**Figure 6.**
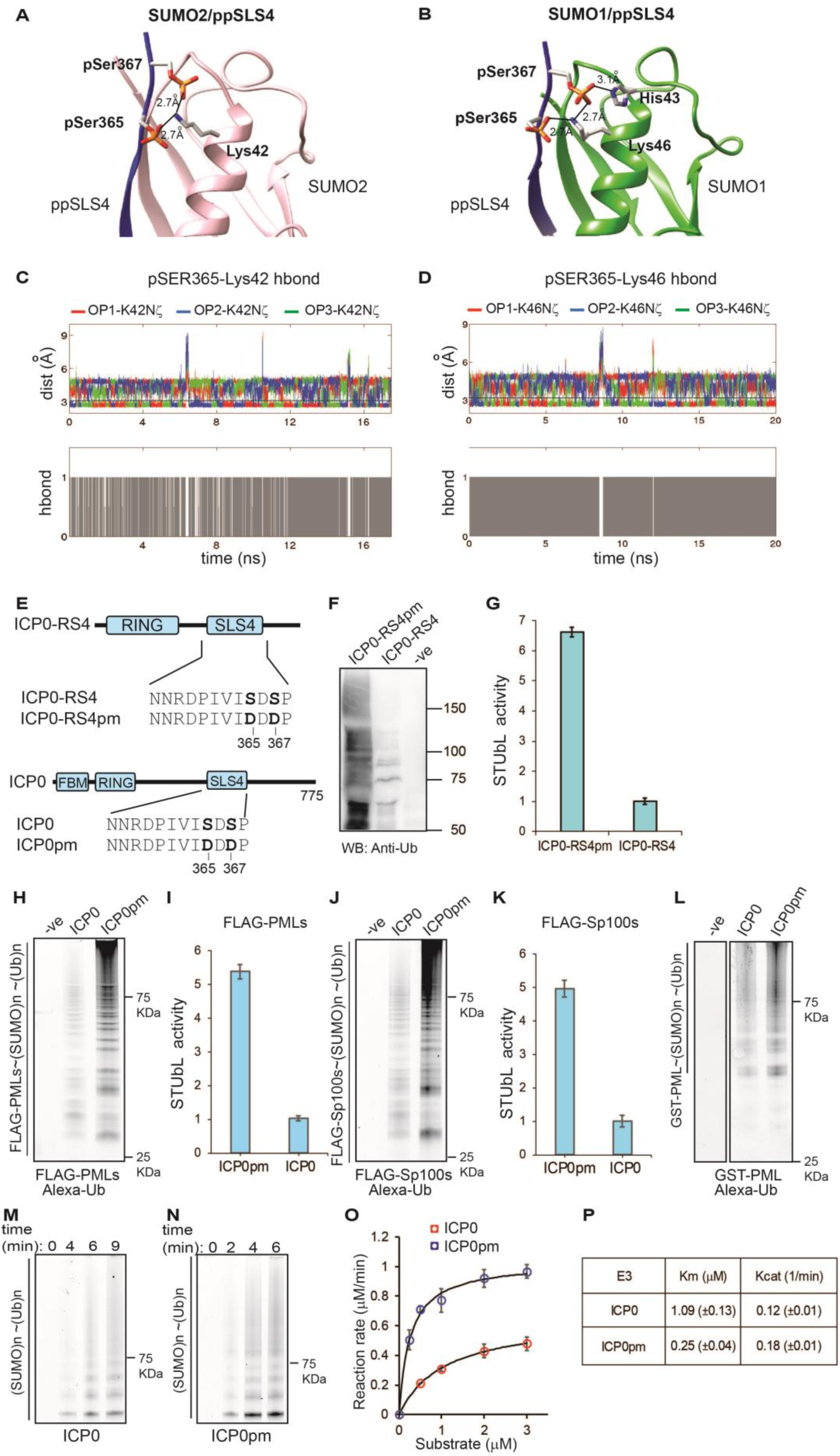
Phosphorylation of SLS4 creates new hydrogen bonds at the SUMO/SLS4 interface to increase the STUbL activity of ICP0. (A) and (B) shows the lowest energy structures of SUMO2/ppSLS4 and SUMO1/ppSLS4 complexes, respectively. The hydrogen bonds between phosphorylated side-chains of SLS4 and SUMO1/2 are shown as black lines. (C) Distances between the three phosphate oxygen atoms of pSer365 and Lysine 42 Nζ of SUMO2 are plotted against time in the MD simulation. Given below is the presence/absence of the hydrogen bond between any one of the phosphorus atoms of pSer365 and Lysine 42 Nζ digitized to a two-stage process (0 or 1), and plotted against time. (D) Similar to (C) plotted for the distance between phosphorylated Ser365 and Lysine 46 Nζ of SUMO1. (E) Schematic of ICP0-RS4, ICP0 and the phosphomimetic version ICP0-RS4pm and ICP0pm. (F) Ubiquitination assays using ICP0-RS4 and the phosphomimetic ICP0-RS4pm as the E3 and SUMO2-chains as the substrate carried out on SUMO2-affinity beads, washed and blotted with the anti-Ub antibody. The (-ve) control lane in this assay lacks the E3. (G) Quantification of the bands observed in (G), where the intensity of ICP0-RS4 lane is normalized to 1. The values are mean of three reactions, and the standard deviation is plotted as error bars. (H) Ubiquitination reaction using SUMOylated PMLs as in Figure 4C, using ICP0 and ICP0pm as the E3 here. (I) The lanes in (H) are quantified and plotted. (J) Same as in (H) using SUMOylated Sp100s as the substrate, and quantified in (K). (L) Same as in Figure 4F, using ICP0 and ICP0pm as the E3. (M) Multi-round ubiquitination reaction as performed in Figure 3I, using 0.5 μM (SUMO)_n_ as the substrate and ICP0 as the E3. The reaction is quenched at different time points (0min, 4min, 6min, and 9min) by adding EDTA and separated on SDS gel. (N) Same as in (M) using ICP0pm as the E3. The time points are (0min, 2min, 4min, and 6min). (O) The reaction rates of ICP0 and ICP0pm are compared at the different substrate ((SUMO2)_n_) concentrations. The values are the mean of triplicates, and the error is the standard deviation of the same. The fit of the data to the Michelis Menten equation is given as black lines. (P) The kinetic parameters of STUbL activity for ICP0 and ICP0pm are given in a table. The values are the mean of triplicates, and the error is the standard deviation of the same.

**Table 2.**
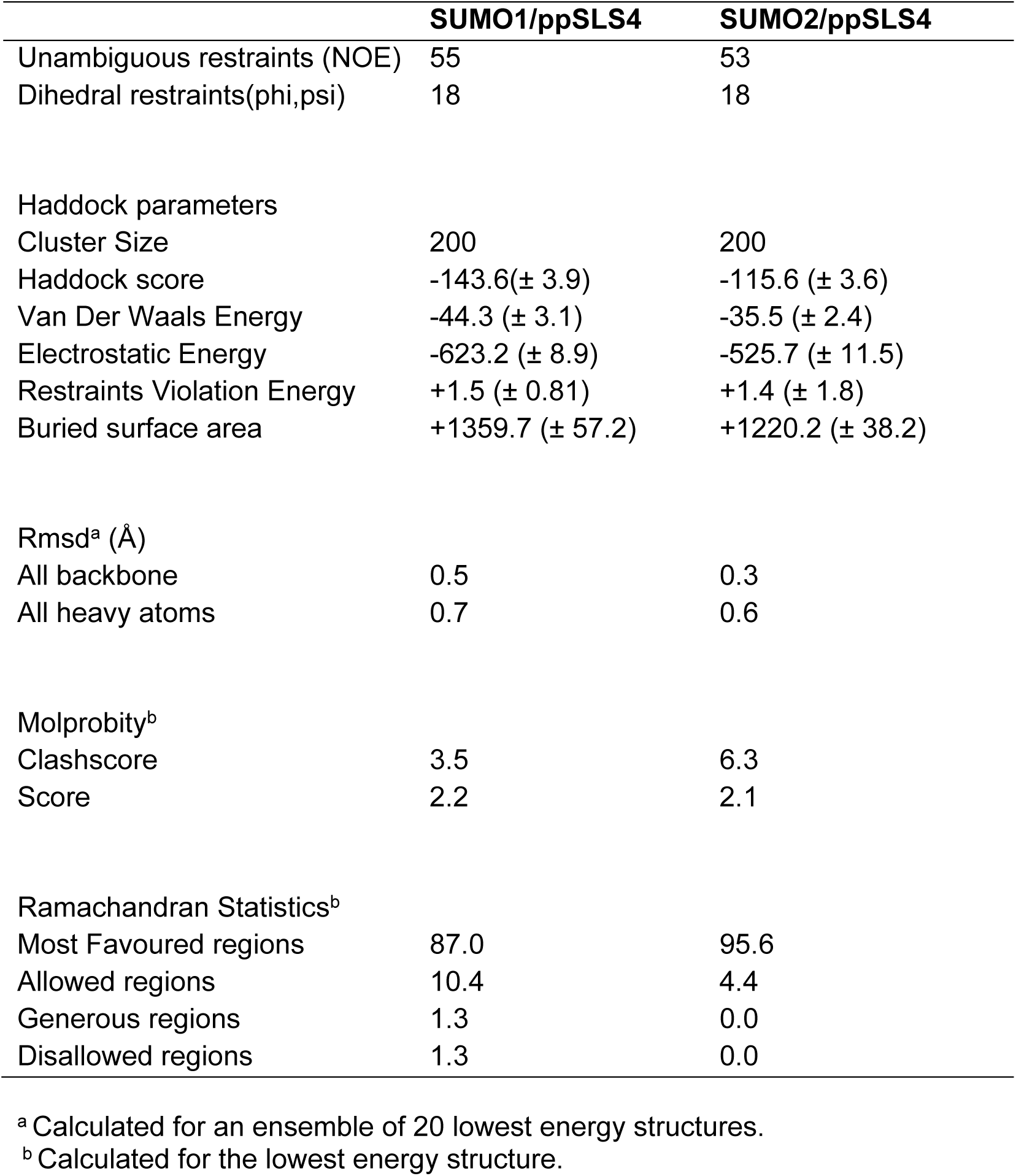
NMR and refinement statistics of the SUMO/ppSLS4 complexes.

The structure of the SUMO1/ppSLS4 complex was studied to determine if a similar mechanism operates between phosphorylated SLS4 and SUMO1. We prepared a sample of ^13^C, ^15^N-labeled SUMO1/ppSLS4 at the stoichiometric ratio of 1:1.5 (SUMO1:ppSLS4). ^13^C,^15^N-filtered (F1), ^13^C,^15^N-edited (F2) NOESY HSQC of this sample detected several intermolecular NOES between SUMO1 and ppSLS4 (Figure S10B), which were used to determine the SUMO1/ppLS4 structure. The twenty lowest energy structures superimposed with a low rmsd of 0.6 Å (Figure S10C). Three new hydrogen bonds were detected in the lowest energy structure (Figure 6B). The oxygen atom of phosphate groups of pSer365 and pSer367 form hydrogen bonds with Lysine 46 present in α1. Phosphorylated Ser367 also forms a hydrogen bond with His43 of SUMO1.

Molecular dynamics simulations of the complexes between phosphorylated SLS4 and SUMO1/2 were carried out to estimate the stability of these new hydrogen bonds observed in the lowest energy structures. The complexes were dissolved in a square water box, equilibrated for 1ns, followed by NVT simulations. The phosphate oxygen atoms of Ser365 and Ser367 formed hydrogen bonds with the Lysine 42 Nζ atom of SUMO2 practically throughout the simulation (Figure 6C). The three oxygen atoms rotated about the central P=O double bond, but at any point in time at least one of them was hydrogen-bonded with Lysine 42. Similarly, the hydrogen bond between Ser367 and Lysine 42 Nζ was found to be reasonably stable (Figure S11A). The hydrogen bonds between phosphoserines and SUMO1 were also stable throughout (Figure 6D and Figure S11B). Collectively, the structures and MD simulations reveal that phosphorylation creates new and stable hydrogen bonds between SUMO and SLS4, leading to the tighter binding of the SUMO/SLS4 complex.

### Phosphorylation increases the STUbL activity of ICP0

The increased affinity between SLS4 and SUMO upon phosphorylation of SLS4 could increase the affinity of ICP0 for SUMOylated substrates. This was tested using a phosphomimetic mutant of ICP0-RS4. Ser365 and Ser367 were mutated to aspartic acids to create a phosphomimetic version of ICP0-RS4 (ICP0-RS4pm, Figure 6E). The same phosphomimetic mutant was also made in the full-length ICP0 (Figure 6E). The ubiquitination activity of ICP0-RS4pm is similar to ICP0-RS4, indicating that phosphorylation does not change the catalytic activity of the RING-finger domain (Figure S6C). Ubiquitination assay was carried out on poly-SUMO2 chains bound to the SUMO-affinity capture beads, washed and blotted with the anti-Ub antibody. In the reaction, the concentration of E3 (ICP0-RS4 or ICP0-RS4pm) was reduced such that the E3: Substrate stoichiometry is ∼1:1. In these conditions, the ICP0-RS4pm had six-fold higher STUbL activity as compared to ICP0-RS4 (Figure 6F and 6G).

The STUbL activity of ICP0 was also compared with its phosphomimetic form ICP0pm, using the SUMOylated PML or Sp100 as the substrates. FLAG-PMLs was SUMOylated, captured on anti-FLAG affinity beads and washed. Subsequently, ubiquitination reaction was carried out on the beads using Ube1 (E1), Ube2d1 (E2), Alexa-Ub, and either ICP0 or ICP0pm as the E3. The reaction was quenched by EDTA after 30min, washed (3x), and the pellet on the beads was separated on SDS gel. The ICP0pm showed a 5-fold increase in STUbL activity compared to the ICP0 (Figure 6H and 6I). When Sp100s was used as the substrate, the effect of phosphorylation was similar (Figure 6J and 6K). The ubiquitination of SUMOylated GST-PML was also enhanced in ICP0pm compared to apo ICP0 (Figure 6L).

Kinetic analysis of the STUbL activity between ICP0 and ICP0pm can determine if phosphorylation increased the intrinsic activity of ICP0 or the affinity towards the substrate. A multi-round ubiquitination kinetics experiments were carried out using either ICP0 or ICP0pm as the E3, and (SUMO2)_n_ chains on SUMO-affinity beads as the substrate. The reactions were quenched with excess EDTA at different time points, washed, separated on SDS gel and quantified (Figure 6M and 6N). The measured ubiquitination rates of ICP0pm were higher than the ICP0 (Figure 6O). A comparison of the kinetic parameters between ICP0 and ICP0pm indicated that phosphorylation did not affect the Kcat of the reaction, but reduced the Km for the substrate by 4-fold (Figure 6P). Altogether, phosphorylation of ICP0 at SLS4 increases its affinity for SUMOylated substrates, and consequently its STUbL.

### Chk2 binds to ICP0 and phosphorylates SLS4

Previous research has indicated that the ATM/Chk2 pathway is important for HSV-1 growth (Li *et al*., 2008). The effect of Chk2 inhibition on HSV1-replication was tested. Vero cells were infected with HSV-1 17 syn+ at the multiplicity of infection (MOI) of 0.1 in the presence and absence of Chk2 inhibitor. Infection in the presence of Chk2 inhibitor exhibit slower progression of Cytopathic Effect (CPE) as compared to the infection in the untreated sample (Figure S12A). HSV-1 mediated cell rounding and detachments were significantly retarded in the presence of Chk2 inhibitor at 36, 48, and 60 hours post-infection (hpi), suggesting that Chk2 inhibition is detrimental to the HSV-1 growth. Virus yield was determined by plaque assay in Vero cells. Chk2 inhibited cells yielded approximately 1000-fold less virus at 48 and 60 hours post-infection (hpi) than the untreated cells, confirming that Chk2 promotes viral growth (Figure 7A).

**Figure 7.**
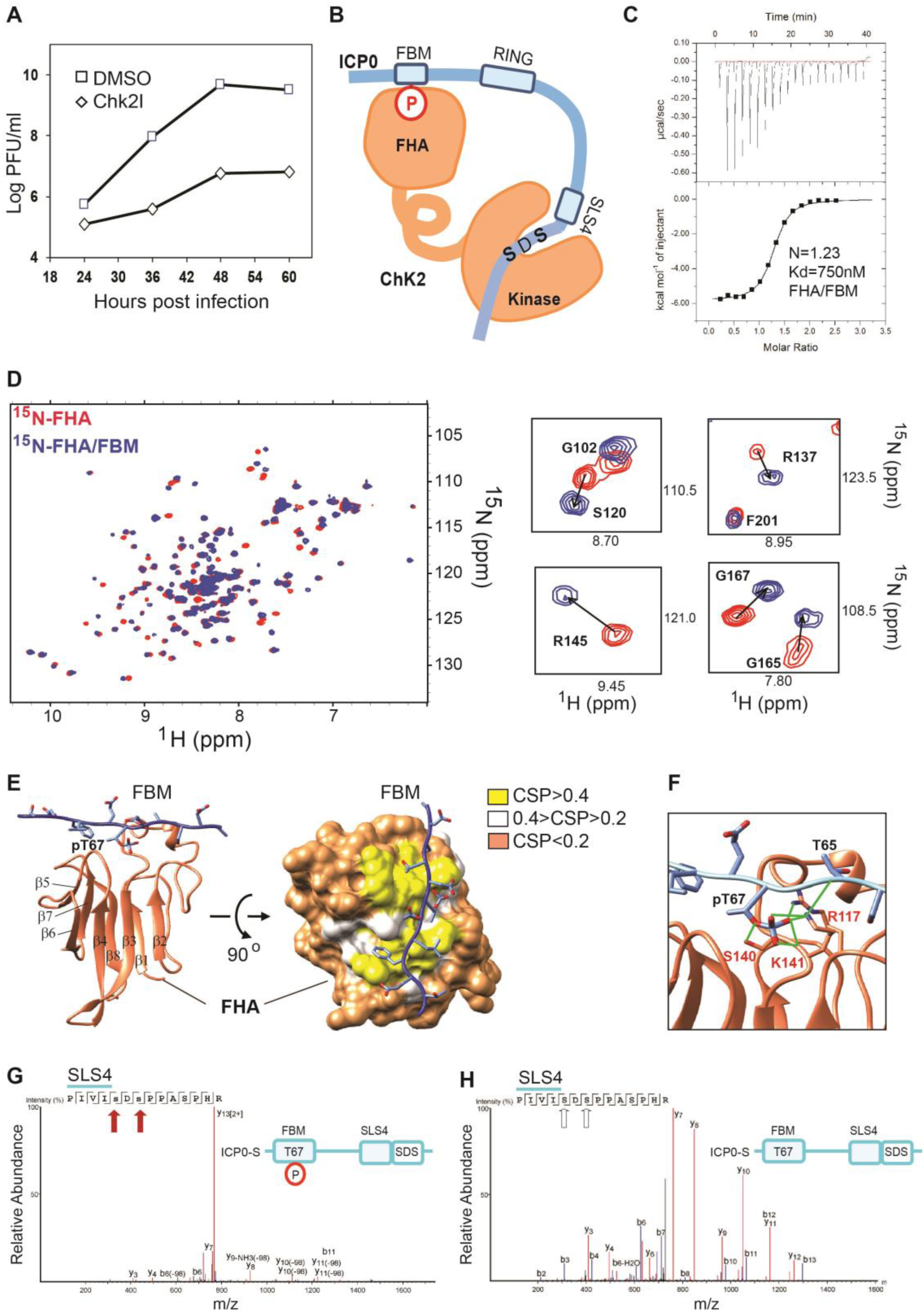
Chk2 interacts with ICP0 and phosphorylates SLS4. (A) Chk2 inhibition suppresses HSV-1 replication. Vero cells were infected at MOI of 0.1 using HSV-1 17 syn+ as described in material and methods. Mock treatment (DMSO) were included as a negative control. Cells and medium were harvested at the indicated time points, and virus yield was assessed by the plaque assay in Vero cells. The values are mean of three experiments. The standard deviation at each time point is minor and hence, plotted separately in Figure S12B. (B) A proposed phosphorylation mechanism of phosphoserines near SLS4, which is dependent on the interaction between Chk2-FHA domain and the phosphorylated FBM from ICP0. (C) ITC binding data of FHA-FBM interaction. The titration curve indicates an exothermic reaction with the dissociation constant Kd = 750 (±50) nM, stoichiometry = 1.23, ΔH = -5.8 (±0.1)kcal/mol and ΔS = 8.5cal/(K.mol.T). (D) An overlay of the ^15^N-edited HSQC of free ^15^N-labeled FHA domain in red and the ^15^N-FHA: FBM (1:1.2) in blue. Expanded regions of the spectra are given on the right. (E) The NMR-data guided HADDOCK structure of the Chk2-FHA/FBM complex. A surface representation of the complex showing observed CSPs on the surface of the FHA domain. (F) The hydrogen bonds observed between FHA and FBM around phosphorylated T67. The hydrogen bonds are colored in green. (G) Schematic of ICP0-S phosphorylated at Thr67, and the MS/MS spectrum of a tryptic peptide of Chk2 treated ICP0-S. The MS/MS data confirms that Chk2 phosphorylates ICP0-S at the phosphoserines S365 and S367 near SLS4 (red arrows). (H) The MS/MS data of Chk2 treated ICP0-S that is not phosphorylated at Thr67 phosphorylation. The serines are not phosphorylated in this case (white arrows).

CK1 phosphorylates ICP0 at Thr67 in the FHA Binding Motif (FBM, aa: 55-74) in ICP0 (Figure 1A). Upon phosphorylation, ICP0 interacts with FHA domains (Chaurushiya *et al*., 2012). Since Chk2 phosphorylates serines, promotes viral replication, and includes an FHA domain, it can potentially bind and phosphorylate ICP0 (Figure 7B). This hypothesis was tested with various tools. The interaction of the phosphorylated FBM with the Chk2-FHA domain was studied by ITC. The binding isotherm given in Figure 7C indicated a strong interaction between FBM and Chk2-FHA. The dissociation constant was measured to be 750nM, consistent with a previous measurement of the FHA/FBM interaction (Chaurushiya *et al*., 2012).

A sample of ^13^C, ^15^N-Chk2-FHA was prepared to obtain molecular details of this interaction by NMR. The backbone chemical shifts of the Chk2-FHA domain were assigned using 3D HNCO, HN(CA)CO, HNCA, and HN(CO)CA experiments (Figure S13A). A synthesized peptide of FBM was titrated into the sample of ^13^C, ^15^N-Chk2-FHA. At 1:0.5 stoichiometric ratio of Chk2-FHA: FBM, the spectra of Chk2-FHA showed a new set of peaks coming from the FBM bound conformation of Chk2-FHA as well as the peaks of the free Chk2-FHA. The simultaneous presence of peaks corresponding to both free and bound conformations indicated that the peaks are in slow exchange compared to the NMR timescale. The slow exchange is a signature of a strong interaction between FBM and the Chk2-FHA, which corroborates the ITC data. At 1:1.2 stoichiometric ratio of Chk2-FHA: FBM, only the peaks corresponding to bound conformation remained in the spectra, implying that Chk2-FHA is saturated with FBM. Figure 7D shows the overlay of free and bound Chk2-FHA. The bound spectra were reassigned using standard 3D NMR experiments. Significant amide chemical shift perturbations were observed between the free and the bound form of the Chk2-FHA domain (Figure S13B). However, Chk2 precipitated at high concentrations (0.2 mM), and further 3D experiments for assigning resonances from side-chains atoms, and intermolecular NOESY experiments were not feasible. Hence, the CSPs were used to dock the FBM to the Chk2-FHA domain (PDB ID: 1GXC) by HADDOCK (Dominguez, Boelens and Bonvin, 2003). Table 3 provides that statistics of the docked structures. Figure 7E shows the lowest energy structure of Chk2-FHA/FBM complex. The surface representation in Figure 7E shows how the CSPs correlate with the FBM interface on the Chk2-FHA.

**Table 3.**
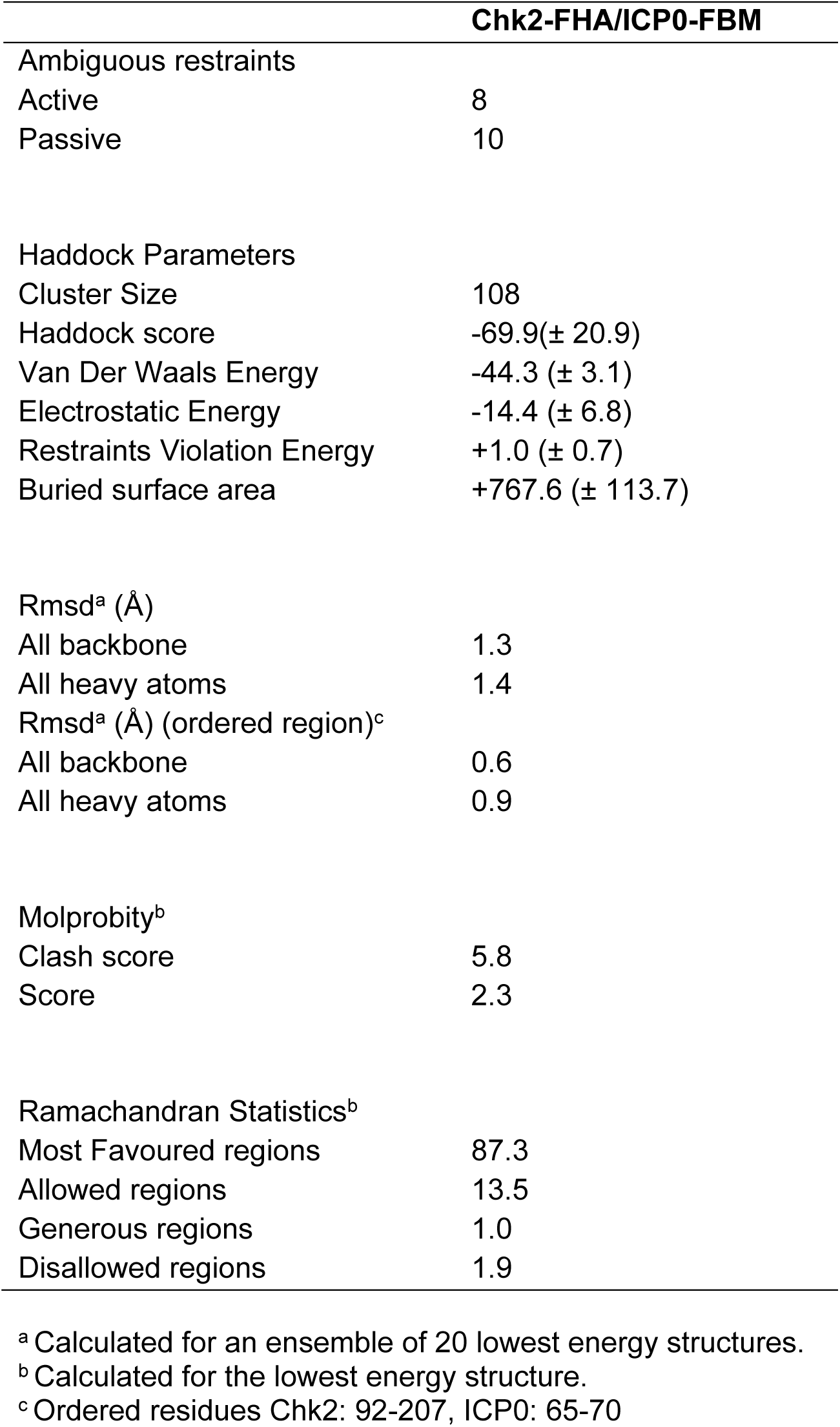
NMR and refinement statistics of the Chk2-FHA/ICP0-FBM complex.

The Chk2-FHA domain has a beta-sandwich fold composed of 8 beta-strands. The N-termini and the C-termini beta-strands align together to form a compact and independent modular domain. The loop between β2 and β3 is extended and also contains a short alpha helix. The phosphorylated FBM binds to the loops between the beta-strands β2/β3, β4/β5, and β7/β8 (Figure 7E). Phosphorylation of Thr67 appears to be essential for this interaction as the phosphate group forms multiple hydrogen bonds with the basic residues in the β2/β3 loop; (i) side-chain of Arg117, (ii) side-chain of Lys141 and (iii) backbone of Lys 141 (Figure 7F). T67 also hydrogen bonds with the side-chain of Ser140 (Figure 7F). Leu193 in the β7/β8 loop forms several hydrophobic contacts with the aromatic ring of Phe70 in FBM (Figure S13C). A couple of the FHA-CHk2 residues interact with the backbone of the FBM and helps to stabilize the polypeptide at the interface. For example, Asn166 in the β4/β5 loop forms two hydrogen bonds with the backbone of Glu68 and Phe70 of FBM (Figure S13C). Similarly, Lys141 side-chain Nζ forms a hydrogen bond with the Thr65 backbone oxygen atom (Figure 7F).

After confirming Chk2-FHA/ICP0-FBM binding, the phosphorylation of ICP0 at SLS4 was checked by MS/MS analysis. A synthetic peptide was designed where the two ICP0 motifs FBM and SLS4 were fused with an intermediate stretch of ∼30 wild-type residues (total 45 residues) to test whether Chk2 can phosphorylate SLS4. Thr67 is phosphorylated in ICP0-S to promote FHA/FBM binding. The chimera is shown in Figure 7G and referred to as ICP0-S. A phosphorylation reaction was initiated with Chk2 as the kinase and the ICP0-S as the substrate. After the reaction, the phosphorylation status of Ser365 and Ser367 were determined by trypsin digestion followed by MS/MS analysis. The mass of peptide fragment corresponding to SLS4 shows that Chk2 phosphorylated SLS4 at both Ser365 and Ser367 (Figure 7G). Some SLS4 was observed where only the Ser367 is phosphorylated (Figure S14), indicating that Chk2 initially phosphorylates Ser367 followed by Ser365. SLS4 phosphorylation was not detected when the ICP0-S is unphosphorylated at Thr67 (Figure 7H). Overall, upon phosphorylation of ICP0 at Thr67 by CK1, Chk2 binds and phosphorylates ICP0 at SLS4.

### CK1 and Chk2 work in tandem to increase the STUbL activity of ICP0

The phosphorylation at SLS4 should increase the STUbL activity of ICP0. The effect of CK1 and Chk2 in the STUbL activity of ICP0 was measured using the substrates used above. Sequential treatment of ICP0 by CK1 and Chk2 increased its STUbL activity for PMLs, Sp100s, and PML (Figure 8A-C). The STUbL activity of CK1 and Chk2 treated ICP0 was similar to the activity of phosphomimetic ICP0pm, indicating that the cumulative effect of CK1 and Chk2 treatment on the STUbL activity is the SLS4 phosphorylation (Figure 8D and 8E). A variant T76E-ICP0 was purified, which mimicked the phosphorylation at T67 by CK1. The STUbL activity of T67E-ICP0 treated with Chk2 alone, compared well with the activity CK1 and Chk2 treated ICP0 for PMLs, Sp100s and PML (Figure 8F-H). The effect of CK1 inhibitors (CK1i) and Chk2 inhibitors (Chk2i) on the STUbL activity of ICP0 were tested. During the phosphorylation reaction, either CK1i or Chk2i were added in the reaction mix. The inhibition of either CK1 or Chk2 depleted the STUbL activity of ICP0 for PMLs, Sp100s and PML (Figure 8I, 8J, and 8K), which indicated that both the kinases were required for SLS4 phosphorylation. However, phosphorylation of ICP0 by CK1 and Chk2 could not rescue the STUbL activity of hydrophobic and charge mutants, suggesting that those residues are central to the SUMO/SLS4 interaction (Figure 8L). Altogether, these results suggest that CK1 and Chk2 act on ICP0 in tandem and enhance its STUbL activity.

**Figure 8.**
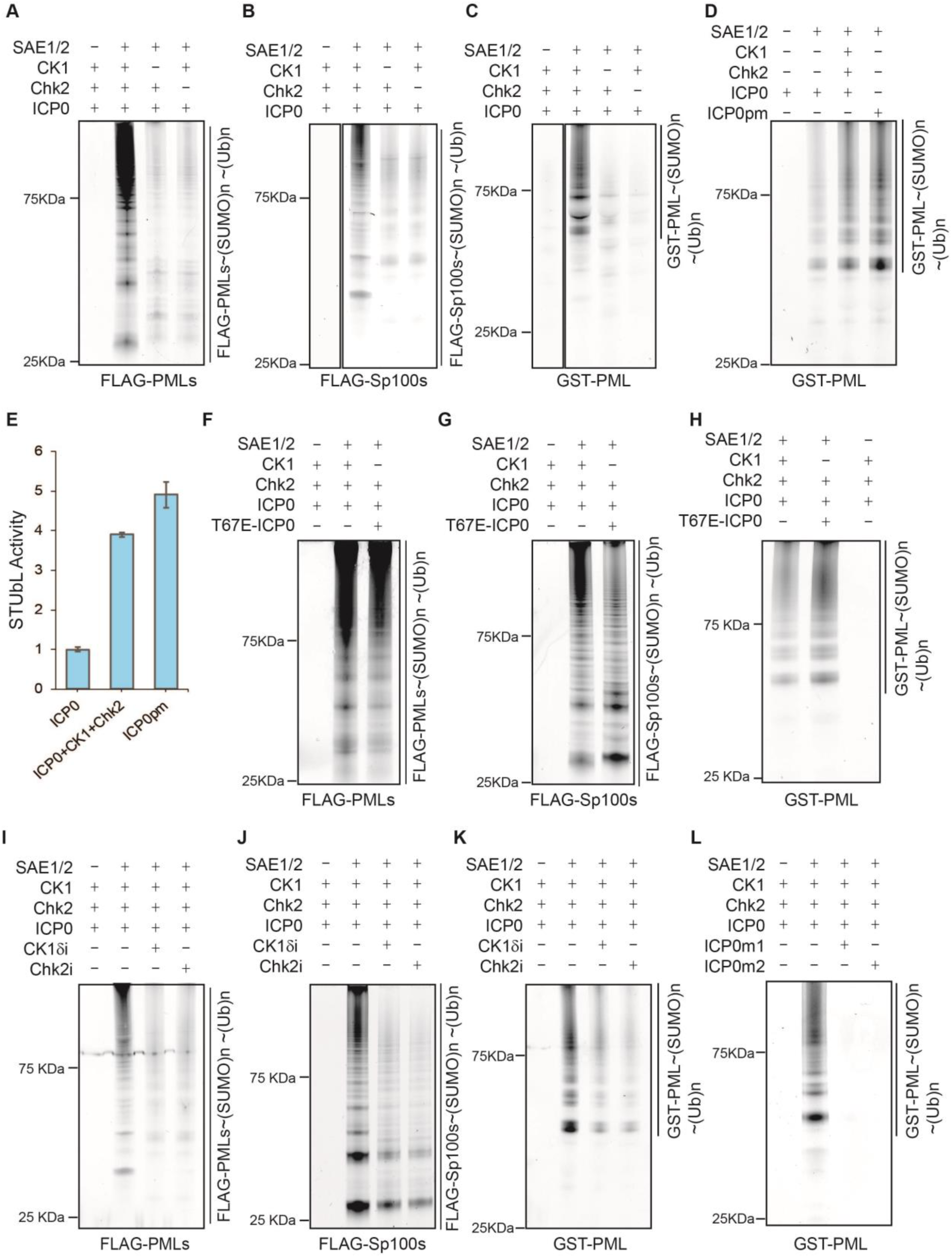
CK1 and Chk2 treated ICP0 has enhanced STUbL activity. Ubiquitination of SUMOylated (A) FLAG-PMLs, (B) FLAG-Sp100s and (C) GST-PML was carried out as in Figure 4, except that ICP0 was pre-treated with either CK1 or Chk2 or both before the start of the ubiquitination reaction. (D) The activation of ICP0 by CK1 and Chk2 was compared with the phosphomimetic mutant ICP0pm, using GST-PML as the substrate. The ubiquitinated bands were quantified and plotted in (E). The values are a mean of three experiments, while the error is the standard deviation of the same. (F) The STUbL activity was compared between CK1+Chk2 treated ICP0, and a phosphomimetic mutant T67E-ICP0, which mimics the phosphorylation of ICP0 by CK1. Here FLAG-PMLs was used as the substrate. FLAG-Sp100s and GST-PML were used as the substrate in (G) and (H). The STUbL activity of CK1 and Chk2 treated ICP0 was compared to its STUbL activity in the presence of either the CK1 inhibitor (CK1i) or the Chk2 inhibitor (Chk2i). ICP0 was treated with CK1, Chk2 and either the CK1i or Chk2i, before using it for ubiquitination of SUMOylated PMLs. The same reactions were also carried out using (J) FLAG-Sp100s and (K) GST-PML as the substrate. (L) The SLS4 mutants ICP0m1 and ICP0m2 were treated with CK1 and Chk2 and used for the ubiquitination of SUMOylated GST-PML. Their activity was compared with CK1, Chk2 treated wt-ICP0. The Ub used in all the assays in this figure is Alexa-Ub and the CK1 isoform used is CK1δ.

## Discussion

ICP0 is transcribed during the immediate early stages of the *lytic* cycle and translocates to the nucleus. There it degrades PML NBs to alleviate host-induced transcriptional repression (Everett, 2006). This process is critically dependent on the STUbL activity of ICP0, by which it can target SUMOylated proteins, assemble polyubiquitin chains on them, and tag them for proteasomal degradation. One or more of the SLS regions in ICP0 are expected to target SUMOylated proteins. Here, a comprehensive study of all the SLS regions identified SLS4 as the only bonafide SIM in ICP0. The structures of SUMO1/SLS4 and SUMO2/SLS4 complexes indicate that both hydrophobic and electrostatic interactions are critical for the SUMO/SLS4 binding. Hydrophobic residues in SUMO1 and SUMO2 create an exposed shallow groove between β2 and α1. The two isoleucines Ile362 and Ile364 in SLS4 bury their side-chains into this groove and forms extensive contacts with the hydrophobic residues of SUMO1 and SUMO2. In addition, the two aspartic acids Asp360 and Asp366 make ionic interactions with the basic residues of β2 and α1. The other SLS sequences do not have the combination of these hydrophobic and acidic residues in the proper arrangement. Hence, they do not bind SUMO1 or SUMO2.

The loss of STUbL activity upon mutations at the interface between SLS4 and SUMO, underline the functional importance of SUMO/SLS4 interaction. Several cellular studies have indicated the importance of the region around SLS4 for degradation of the PML NBs (Boutell *et al*., 2011; Everett *et al*., 2014b; Muller, 2014; Zheng and Gu, 2015; Zheng, Samrat and Gu, 2016). Intriguingly, the C-terminal region, including SLS5-7 is also important for PML body degradation (Gu and Zheng, 2013). However, the NMR titrations showed that individually SLS5, SLS6, and SLS7 do not bind SUMO. Moreover, the entire region of SLS5-7 also did not effectively interact with SUMO. A recent study postulated that SLS5-7 might be a portion of a larger structural fold (Everett *et al*., 2014b). Our results strengthen the possibility that SLS5-7 constitutes a structural fold that either binds SUMO or other PML body constituent proteins via a mechanism that is entirely different from the typical SUMO/SIM interaction.

Phosphorylation of viral proteins is important for *lytic* reactivation in the *herpesviridae* family (Yue, Gershburg and Pagano, 2005; El-Guindy *et al*., 2007; Kenney and Mertz, 2014; Purushothaman, Uppal and Verma, 2015). Phosphorylation of ICP0 is important for PML NBs disruption (Boutell *et al*., 2008; Mostafa *et al*., 2011). Specifically, HSV-1 is defective in neuronal replication and explants-induced reactivation from latency when Ser365, Ser367, and Ser371 are mutated to alanines in ICP0 (Mostafa *et al*., 2011). The mutant was unable to dissociate and degrade SUMOylated PML from the PML NBs (Boutell *et al*., 2008). Ser371 is far from the binding interface to create an impact on the SUMO/SLS4 binding. However, we show that phosphorylation of Ser365 and Ser367 dramatically increased the SUMO/SLS4 affinity by ∼25 fold. The higher affinity translates into enhanced ubiquitination of SUMOylated substrates, indicating that phosphorylation of ICP0 at SLS4 can switch it into a potent STUbL. PML NBs are short-lived post association with ICP0. Though SUMO/SLS4 affinity is weak, a couple of factors must help ICP0 to degrade SUMOylated proteins in PML NBs rapidly. First, the constitutive proteins in PML NBs like PML, Sp100, and hDaxx are heavily SUMOylated, leading to a high local concentration of SUMO in the PML NBs. Hence, the effective affinity due to local concentration will be higher than the *in-vitro* measured affinity. Second, phosphorylation at SLS4 increases the affinity towards SUMO, which enhances the STUbL activity. ICP0 that cannot be phosphorylated at SLS4 is inactive in degrading PML NBs (Boutell *et al*., 2008). Hence, the switch in STUbL activity upon phosphorylation could be essential to swiftly counter the emerging PML NBs as the viral genome enters the nucleus.

Viral replication exploits several host signaling pathways; however, the molecular mechanism of this process is often ambiguous. The HSV-1 directly activates and exploits the ATM/Chk2 pathway for an effective infection (Everett, 2006). Since these pathways overlap with antiviral responses, it seems counterintuitive as to why HSV-1 would activate these pathways. This work provides a clue to resolve this conundrum by showing that ICP0 exploits the activated Chk2 to phosphorylate SLS4, and then efficiently ubiquitinates SUMOylated nuclear proteins to tag them for degradation (Figure 9). The SLS4 region is highly charged due to the presence of multiple negatively charged residues, and due to technical difficulties with quantitative mass-spectrometry of charged peptide fragments, we could not confirm by Chk2 inhibition experiments, whether ChK2 is the sole kinase that phosphorylates SLS4 *in vivo*. However, multiple results implicate Chk2 as the relevant kinase for ICP0 (Figure 7). Inhibition of Chk2 significantly retarded CPE progression of HSV-1 and reduced the viral growth by 1000-fold. *In-vitro* phosphorylation reactions revealed that Chk2 specifically targets Ser365 and Ser367 at SLS4. The phosphorylation by Chk2 is, however, dependent on phosphorylation by CK1. These two kinases work in tandem to activate ICP0 into a potent STUbL during the *lytic* reactivation.

**Figure 9.**
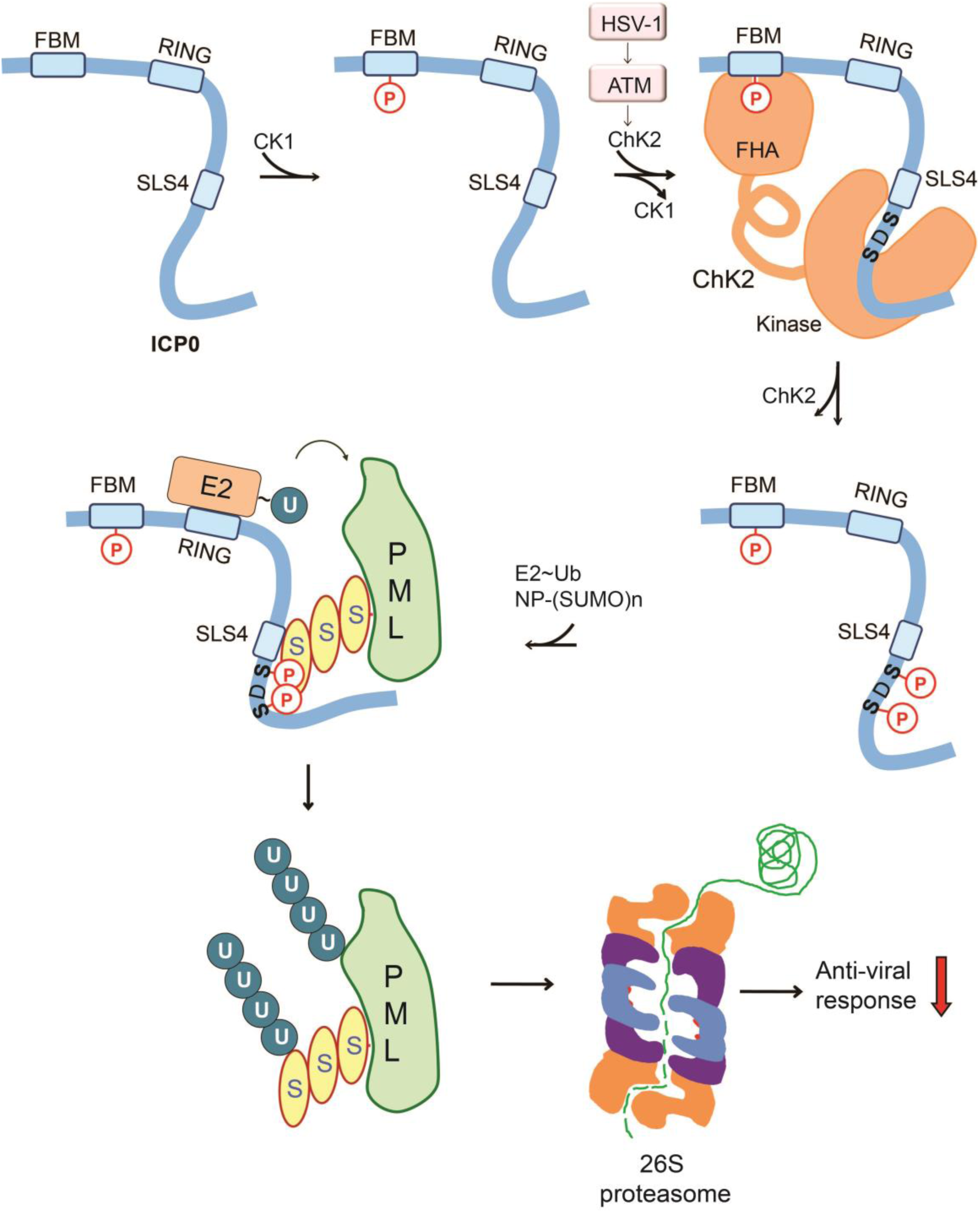
The proposed model for Chk2-mediated activation of ICP0. CK1 phosphorylates the N-terminal FBM at Thr67. HSV-1 activates ATM/Chk2 pathway, and the activated Chk2 binds ICP0 to phosphorylate the phosphoserines at SLS4. The phosphorylated ICP0 identifies SUMOylated nuclear proteins like PML with higher affinity and assembles polyubiquitin chains using the E2∼Ub to tag them for degradation by the 26S proteasome.

Crosstalk between PTMs is a known trend in eukaryotic biology (Hunter, 2007). For example, phosphorylation can increase as well as inhibit ubiquitination (Sheng *et al*., 2002; Hunter, 2007; Dou *et al*., 2012). SUMOylation and ubiquitination can compete for the same lysine in a protein (Hoege *et al*., 2002). Phosphorylation can promote SUMOylation through the phosphorylation-dependent SUMOylation motif (Hietakangas *et al*., 2006). In some cases, phosphorylation can also negatively impact SUMOylation as observed for c-Myb (Bies, Sramko and Wolff, 2013). Until now, these three PTMs were found to cross-talk in a pair-wise manner. In this study, we find that phosphorylation increases the activity of a ubiquitin ligase in a SUMO-dependent manner (Figure 9). To our knowledge, this is the first instance where all the three PTMs ubiquitination, SUMOylation, and phosphorylation are simultaneously involved in a cross-talk. This observation opens up discussions on how the virus turns host defenses against the host by using multiple PTMs. SIMs in several other STUbLs such as Uls1, Slx5, Slx8 (*S. Cerevisiae*), Rfp1 (*S. pombe*), and, RNF4, Rad18, and Arkadia (*H. Sapiens*) have phosphoserines nearby (Prudden *et al*., 2007; Sriramachandran and Dohmen, 2014). It remains to be seen whether phosphorylation regulated STUbL activity is present in host E3s or is exclusive to the viral E3s.

It is rather intriguing that HSV-1 has evolved to use a complex mechanism of two kinases to activate ICP0. As a part of the antiviral response, rescue factors like RNF8 and RNF168 assemble at the viral replication nodes to inhibit transcription of the viral genome. Weitzman and co-workers have demonstrated that upon phosphorylation by CK1, ICP0 binds to RNF8-FHA domain, and degrades RNF8 (Chaurushiya *et al*., 2012). Our results show that subsequent to CK1, Chk2 phosphorylates and activates ICP0 to degrade SUMOylated PML NB proteins. Hence, the phosphorylation of FBM motif ensures that inhibition due to both the RNF8 and the PML NBs are efficiently countered. Interestingly, while RNF8 degradation occurs in a SUMO-independent manner, degradation of PML NBs occurs in a SUMO-dependent mechanism. Given both the host responses (RNF8 and PML NBs) are negated by the FHA/FBM interaction, this interaction now becomes an interesting target for therapeutic intervention. The mechanism of using two kinases could be necessary to coordinate and regulate the activity of ICP0 at various stages of infection. Evidently, the viral E3 ICP0 has developed a meticulous strategy to modulate the host antiviral responses for efficient viral replication. We find that the cross-talk between multiple PTMs plays a vital role in its strategy.

### Accession Numbers

The coordinates of SUMO2/SLS4 and SUMO1/SLS4 are deposited in the PDB under accession codes 5H4X and 5H4T, respectively. The coordinates of SUMO2/ppSLS4 and SUMO1/ppSLS4 were deposited in the PDB under accession codes 5H51 and 5H4W, respectively. The backbone assignments of the Chk2 FHA domain were deposited in BMRB under accession code 27042.

## Acknowledgments

The NMR data were acquired at the NCBS-TIFR NMR Facility. The authors would like to thank Purushotham Reddy for helping with NMR data acquisition. The MS data were collected at VProteomics (New Delhi) and at NCBS Proteomics facility. The SUMO1 plasmid was a gift from Rama Koti (TIFR). Chk2-FHA plasmid was kindly provided by Ashok Venkitaraman (InStem). R. Andrew Byrd (NCI) and Allan Weissman (NCI) are acknowledged for critical comments on the manuscript. H. N is supported by a fellowship from CSIR-UGC, Government of India. R.D. is the recipient of Ramalingaswamy fellowship from Department of Biotechnology (DBT), Government of India. This research was funded by intramural grants from the National Center for Biological Sciences, Tata Institute of Fundamental Research.

## Materials and Methods

### Plasmids and Peptides

Synthetic genes for SUMO2, ICP0-R, and ICP0-RS4, PML were obtained from Lifetech, which were subsequently cloned in expression vectors. SUMO2 was cloned into pET3a between NdeI and BamHI restriction sites, while ICP0-R, ICP0-RS4, and PML were cloned into pGEX6P-1 between BamHI and NotI sites. The pQE80L-SUMO1 clone was obtained from Dr. Koti, TIFR, Mumbai. Ube2d1, SUMO-E1, and UBC9 (SUMO-E2) constructs were obtained from Addgene. Chk2-FHA plasmid was a gift from Ashok Venkitaraman (MRC), and Ube1 plasmid was a gift from Cynthia Wolberger (JHU). SENP2 was a gift from Chris Lima (MSKCC). Full-length ICP0 was synthesized in a pFastHtb vector (Invitrogen). Mutants of ICP0 was made by site-directed mutagenesis. All the synthetic peptides were purchased from Lifetein LLC as lyophilized powders.

### Protein Expression and Purification

All the proteins were expressed in BL21 DE3 either in Luria broth for unlabelled samples or in ^15^NH_4_Cl/^13^C_6_-glucose M9 media for labeled samples. SUMO1, SUMO2, SUMO-E1, and SUMO-E2 (UBC9) were expressed and purified as discussed earlier (Chatterjee, Tripathi and Das, 2019). Ube2d1 was purified similar to UBC9. Ube1 and Ub are purified, as discussed earlier (Das *et al*., 2013).

GST-ICP0-R and GST-ICP0-RS4 were cultured at 37°C to 0.6 OD_600_ and were induced at 18°C with 1mM IPTG. Cells were lysed by sonication in PBS (supplemented with protease inhibitors). The lysate was clarified by centrifugation and supernatant was passes through pre-equilibrated Glutathione Agarose (Protino ®) beads for binding. The binding was followed by a high salt wash with 0.5 M NaCl in 50mM Phosphate, 0.1% Triton X-100 and 0.05% sodium azide. Next, the beads were washed with PBS, 0.1% Triton X-100, and 0.05% sodium azide. The protein was eluted with 20mM reduced glutathione in PBS buffer. Glutathione was subsequently removed by dialysis. GST-PML was purified similar to ICP0-R, except a last step of purification by gel-filtration was carried out in PBS.

M9 culture of Chk2-FHA (aa:64-212) was induced at 18°C for 16 hours. The culture was harvested, and cells were resuspended in 50mM phosphate buffer with 300mM NaCl followed by sonication. The supernatant was bound to Ni^2+^ NTA-agarose beads (Protino), washed with 50mM phosphate, 300mM NaCl, and 20mM Imidazole buffer. The protein was eluted with 50mM phosphate, 300mM NaCl with a gradient of 100∼200mM Imidazole. The purified protein was injected into a gel filtration (Superdex75 16/600 column) equilibrated with 50mM phosphate, 300mM NaCl, and 2mM DTT at pH 7.4.

To produce full-length ICP0, *Sf*9 cells were grown in 1X 10 ^6^ for 1 litre of Sf900 II (Gibco) media in presence of antibiotic and antimycotic (Gibco) at 27 °C for 3 days. 50 ml of P3 was added to the *Sf*9 cells. The cells were incubated for another three days, after which, the cells were harvested and lysed in 50mM Tris-Cl (pH 8.0), 500 mM NaCl, 0.01 % tween 20, 2mM EDTA protease inhibitor (Roche), 100 mM PMSF. The supernatant after centrifugation was purified in by Ni-NTA purification. The final samples was dialysed in 50mM Tris-HCl (pH 8.0), 150 mM NaCl and 2mM EDTA protease inhibitor (Roche). The mutants of ICP0, were grown and purified in the similar manner as the wt-ICP0.

### NMR experiments

The NMR spectra of SUMO1 and SUMO2 were recorded at 298K on 800 MHz Bruker Avance III HD spectrometer with a cryoprobe head, processed with NMRpipe(Delaglio *et al*., 1995) and analyzed with Sparky(Kneller and Kuntz, 1993). The SUMO and SUMO/SLS4 NMR samples were prepared in PBS buffer, with 5mM DTT at pH 7.4 and 10% D_2_O. Standard CBCA(CO)NH, HNCACB, HNCA, HN(CO)CA, HNCO and HN(CA)CO experiments were used for backbone assignments and HCCH-TOCSY for side-chain assignments of the SUMO1 and SUMO2. ^1^H-^1^H TOCSY and ^1^H-^1^H NOESY were acquired and used to assign SLS4. For NMR titration experiments, ∼3 mM peptides were titrated into ∼0.5 mM ^15^N-SUMO1/2. The titration data was fit in 1:1 protein: ligand model using the equation CSP_obs_ = CSP_max_ {([P]_t_+[L]_t_+K_d_) - [([P]_t_+[L]_t_+K_d_)^2^-4[P]_t_[L]_t_]^1/2^}/2[P]_t_, where [P]_t_ and [L]_t_ are total concentrations of protein and ligand at any titration point. Intermolecular NOEs between SUMO1 and SLS4 were obtained by acquiring a ^15^N-edited NOESY-HSQC on a ^2^H, ^15^N-SUMO1/SLS4 (1:5) complex sample with a mixing time of 300 ms. A reference ^15^N-edited NOESY-HSQC of free SUMO1 was used to rule out any intramolecular NOEs arising from incomplete deuteration of SUMO1. Intermolecular NOEs between SUMO2 and SLS4 were measured similarly. ^13^C,^15^N-filtered (F1), ^13^C,^15^N-edited (F2) NOESY HSQC was collected on a ^13^C, ^15^N-SUMO1/ppSLS4 (1:1.5) complex sample with a mixing time of 200ms to measure intermolecular NOEs between SUMO1 and ppSLS4. A similar experiment was performed to obtain intermolecular NOEs in the SUMO2/ppSLS4 complex. The Chk2-FHA domain was assigned by standard HNCO, HN(CA)CO, HNCA, HN(CO)CA experiments on a 180μM ^13^C-^15^N-FHA-Chk2 sample. For NMR titration experiments, 2.4mM ICP0-FBM was titrated into 65 μM ^15^N-labeled Chk2-FHA domain.

### Structure Calculations

Unambiguous restraints between the SUMO1 and SLS4 were determined from the intermolecular observed NOEs. The assignments of SLS4 resonances were carried out by 2D ^1^H-^1^H TOCSY and ^1^H-^1^H NOESY experiments. The extended structure of SLS4 was determined using torsion angles from ^1^Hα assignments and ^1^H-^1^H NOESY restraints in xplor-NIH. The solution structure of SUM01/SLS4 complex was calculated in HADDOCK(Dominguez, Boelens and Bonvin, 2003) using the structure of SUMO1 (PDB ID: 4WJO) and the extended structure of SLS4. Rigid body energy minimization generated one thousand initial complex structures, and the best 200 lowest energy structures were selected for torsion angle dynamics and subsequent Cartesian dynamics in an explicit water solvent.

Default scaling for energy terms was applied. The interface of SUMO1 was kept semi-flexible during simulated annealing and the water refinement steps. Following the standard benchmarked protocol, cluster analysis of the 200 water-refined structures yielded a single clear ensemble. The SUMO2/SLS4 complex was similarly docked, except that intermolecular restraints between SUMO2 and SLS4 were used and the starting structure of SUMO2 was taken from a crystal structure of free SUMO2 (PDB ID: 1WM3). The SUMO1/ppSLS4 and SUMO2/ppSLS4 complexes were docked using intermolecular NOES obtained from the ^13^C,^15^N-filtered (F1), ^13^C,^15^N-edited (F2) NOESY HSQC experiments. An extended structure FBM peptide was obtained in the manner similar to SLS4. We obtained the Chk2-FHA/FBM model structure by docking the Chk2-FHA (PDB ID: 1GXC) and the FBM peptide using chemical shift perturbations observed in the NMR titration experiments.

### *In vitro* assays

The *in-vitro* polyubiquitination assays were performed using Ube1 (0.5 μM), Ube2d1 (3 μM), E3s ICP0-R, ICP0-RS4 and its mutants (5 μM) and 50 μM Ub. The reaction mixture was incubated at 37°C for 45 min in the reaction buffer containing 50 mM Tris pH 7.5, 50 mM NaCl, 2 mM MgCl_2,_ and 5 mM ATP. The reaction was quenched with 5 mM EDTA, 4xSDS gel loading buffer, and resolved on 12% Bis-Tris gel. The gel was transferred onto PVDF and blotted with anti-Ub antibody (ENZO-P4D1) using manufacturer protocol. The chemiluminescence was observed by clarity ECL (Bio-Rad) staining in Image Quant LAS 4000 (GE). The SUMO2-chains were used as a substrate (100 ng) and blotted with the anti-SUMO2 antibody to observe ubiquitination of SUMO2-chains. In another set of reactions, the ubiquitination was carried out on SUMO-capture affinity beads (Enzo), quenched and washed (3x) with high salt-buffer, followed by blotting using anti-Ub monoclonal antibody.

For the assay with a phosphomimetic mutant of ICP0-RS4, the concentration of ICP0-RS4 and ICP0-RS4pm was 0.5 μM, and the reaction time was 20mins. The STUbL assays with full-length ICP0 (or its mutants) with SUMO2-chains were done by incubating Ube1 (1 μM), Ube2d1 (5 μM), ICP0 or its mutants (10 μM), Alexa Fluor Maleimide (Invitrogen) labeled Ub (20 μM) with SUMO2-chains on SUMO-capture beads for 30 min at 37°C, quenched and washed (3x), and separated on a 12% Bis-Tris SDS gel and visualized in Uvitec (Cambridge). For the STUbL assays with FLAG-PMLs or FLAG-Sp100s, a SUMOylation reaction was first carried out with SAE1/2 (4 μM), Ubc9 (30 μM), SUMO2 (150 μM) and FLAG-PMLs or FLAG-Sp100s (150 μM) at 37°C. The reaction was subsequently incubated on anti-FLAG affinity beads (Sigma), washed (3x) and the ubiquitination assay was carried out on the beads similar to that for SUMO2-chains. The STUbL assays with GST-PML were performed similarly, except the reaction was incubated on Protino Glutathione agarose (Thermo Scientific). For the kinetic analysis with ICP0 and ICP0pm, the enzyme concentrations were Ube1 (1 μM), Ube2d1 (2 μM), ICP0 or ICP0pm (2 μM) and SUMO2-chains (0-3 μM). The deSUMOylation reactions were carried out by treating with 5 μM SENP2 for 2-3 hours.

For the phosphorylation assays, 10 μg ICP0-S was incubated with 0.5 μg Chk2 (Origene) for 30 min at 30°C in a 20 μl reaction containing 50 mM Tris (pH 7.5), 10 mM MgCl_2_, 10 mM MnCl_2_, 5 mM DTT and 5 mM ATP. The entire phosphorylation reaction was directly digested with trypsin and used for mass spectrometry. Phosphorylation assays of ICP0 with CK1 and Chk2, were performed by incubating 20 μg his_6_-ICP0 with 0.5 μg CK1δ (Merck Millipore) and 0.5 μg ChK2 (Origene) in buffer containing 50 mM Tris (pH 7.5), 10 mM MgCl_2_, 10 mM MnCl_2_, 5 mM DTT and 5 mM ATP on Ni^2+^ NTA-agarose beads (Protino). Subsequent to the kinase reaction, beads were washed (3x), and ICP0 was eluted by 200 mM Imidazole, dialyzed and used for STUbL assays. CK1 inhibition was carried out by using PF5006739 (Sigma) at 0.2 mM concentration. ChK2 inhibition was carried out by using 2-(4-(4-Chlorophenoxy) phenyl)-1H-benzimidazole-5-carboxamide hydrate (Sigma) at 0.2 mM concentration.

### Cell culture and transfection

HEK293T cells were maintained in DMEM with 10% serum. One 100mm dish was transfected with 10µg GFP-ICP0 plasmid with Lipofectamine 3000. Immunoprecipitation (IP) was performed by 36-hours post-transfection. Anti-GFP sepharose beads (CST) were used for IP, according to the protocol provided by the manufacturer. After IP, beads were directly loaded onto reducing SDS PAGE to resolve immunoprecipitated proteins. The band, matching the size of the protein of interest, was excised and analyzed by Mass Spectrometry

### Mass-spectrometry

Samples were digested in the NCBS proteomics facility using an established protocol (Wiese *et al*., 2007). For LC-LTQ Orbitrap MS analysis, samples were resolubilized in 2% [v/v] acetonitrile, 0.1% [v/v] formic acid in water and injected onto an Agilent1200 (Agilent, Santa Clara, CA, USA) nano-flow LC system that was in-line coupled to the nano-electrospray source of a LTQ-Orbitrap Discovery hybrid mass spectrometer (Thermo Scientific, SanJose, CA, USA). Peptides were separated on Zorbax 300SB-C18 (Agilent, Santa Clara, CA, USA) by a gradient developed from 2% [v/v] acetonitrile, 0.1% [v/v] formic acid to 80% [v/v] acetonitrile, 0.1% [v/v] formic acid in water over 70 min at a flow-rate of 300 ml/min. Full MS in a mass range between m/z 300 and m/z 2000 was performed on an Orbitrap mass analyzer with are a solution of 30,000 at m/z 400 and an AGC target of 2 X10^5^. The strongest five signals were selected for CID–MS/MS in the LTQ ion trap at a normalized collision energy of 35% using an AGC target of 1 x 10^5^ and two micro scans. Dynamic exclusion was enabled with one repeat counts during 45s and an exclusion period of 120s.

Peptide identification was performed by CID-based MS/MS of the selected precursors, which also revealed the site of phosphorylation modification. For protein/peptide identification, MS/MS data were searched against the custom peptide sequence provided by Lifetein using PEAKS Studio X software. The search was set up for full tryptic peptides with a maximum of three missed cleavage sites. Oxidized methionine, serine, and threonine were included as variable modifications. The precursor mass tolerance threshold was 10ppm, and the maximum fragment mass error was 0.5 Da. The peptide identification was made at 1 % FDR.

### Viral growth and plaque assay

The Vero cell line was purchased from NCCS Pune cell repository and was grown at 37°C, and 5% CO2 in Dulbecco’s modified Eagle’s medium (DMEM) supplemented with 10% FBS, 100 U/ml of penicillin and 100 μg/ml of streptomycin. The wt-HSV1 17 syn+ strain was a kind gift from Karen L Mossman, which was propagated and tittered in Vero cell line (Brown, Ritchie and Subak-Sharpe, 1973). Vero cells were grown in a 60 mm tissue culture dish. Before infection, the cells were pre-treated with CHK2 inhibitor (10 µM) for 1 hour. Control cells were treated with DMSO. Infection was carried out using wt-HSV1 17 syn+ strain at MOI of 0.1 for two hours at 37°C. After two-hour viral inoculum was removed and fresh medium was added with inhibitor. Cells and medium were harvested at various time point post-infection (12, 24, 36, 48 and 60 hours), then lysed by freeze and thaw cycles. Serial ten-fold dilutions of the lysate were prepared. Each dilution was used to infect the monolayers of Vero cell lines in triplicate. Infection was carried out for two hours at 37°C with intermittent rocking. Post-infection, the viral inoculum was removed, and fresh DMEM medium containing 1% heat-inactivated serum and 0.5% methylcellulose was added. After three days incubation at 37°C, cells were washed with PBS and fixed with methanol for 15 minutes at room temperature followed by staining with 1X Giemsa stain. The plaques were counted and imaged using a 4X objective in a fluorescence microscope. The proteomics data is submitted at PRIDE with accession number PXD014671.

### Molecular Dynamics simulations

All-atom MD simulations of SUMO1/ppSLS4 and SUMO2/ppSLS4 were run on GPU clusters using NAMD (Phillips *et al*., 2005). The protein and ions were described with the CHARMM36 force field (MacKerell, Feig and Brooks, 2004; Huang and Mackerell, 2013) and water molecules with the TIP3P model. The proteins were solvated in water box extending 12 Å from the outermost protein atom. The simulations were started from experimentally determined NMR structures and were energy minimized. The energy-minimized structures were allowed to equilibrate for 5 ns before the production run started. A time step of 2fs was used with the bonds involving hydrogen atoms being constrained using the SHAKE algorithm (Ryckaert, Ciccotti and Berendsen, 1977; Ryckaert, 1985). Electrostatic interactions were calculated using the PME method (Essmann *et al*., 1995), and the van der Waals interactions were truncated beyond 12 Å. Periodic boundary conditions were imposed in all directions. The temperature of the systems was controlled at 300 K using the Langevin dynamics, and the pressure was kept at 1 atm using the Nose-Hoover Langevin piston method (Martyna, Tobias and Klein, 1994; Feller *et al*., 1995). The data were analyzed and plotted using VMD (Humphrey, Dalke and Schulten, 1996), UCSF-Chimera (Pettersen *et al*., 2004), and Matlab.

### Isothermal Titration Calorimetry

The purified FHA-Chk2 and the synthetic peptide of ICP0-FBM were dissolved in buffer (50 mM phosphate, 300 mM NaCl pH 7.4). The 750 µM peptide solution was injected into a sample cell containing 50 µM FHA-Chk2 at 25°C. The measurement was performed on a Microcal ITC_200_ (GE Healthcare), and binding isotherm was plotted and analyzed using Origin (v7.0). Integrated interaction heat values from individual experiments were normalized, and the data were fit, omitting the first point, using a single binding site model.

