## Supplemental Information for "The viral SUMO-targeted Ubiquitin Ligase ICP0 is phosphorylated and activated by host kinase Chk2"

### Supplementary Information

Dambarudhar SS Hembram[1], Hitendra Negi[1,2], Poulomi Biswas[1,3], Vasvi Tripathi[1], Lokesh Bhushan[1], Divya Shet[1,4], Vikas Kumar[1,5] and Ranabir Das[1*]

[1]National Center for Biological Sciences, TIFR, Bangalore, India

[2]SASTRA University, Thirumalaisamudram, Thanjavur, India

[3]Current address: Indian Institute of Technology, Kharagpur.

[4]Current address: Raman Research Institute, Bangalore.

[5]Current address: University of Nebraska Medical Center, Omaha, NE.

*Electronic address:

List of Contents — Page


1. Supplementary Table S1 — 3
2. Supplementary Table S2 — 4
3. Supplementary Table S3 — 4
4. Supplementary Figures S1-S11 — 5-22

**Table S1.** Peptide sequences used to study the interaction between SUMO1/2 and SIM Like Sequences (SLS) in ICP0

| SLS | Peptide Sequence |
|---|---|
| SLS-1 (163-167) | $^{156}$NAKLVY***LIV***G***V***TPSGS$^{172}$ |
| SLS-2 (176-179) | $^{170}$SGSFST***IPIV***NDPQTRMEAEE$^{190}$ |
| SLS-3 (331-334) | $^{325}$GGVGVG***VGVV***EAEAGRPR$^{342}$ |
| SLS-4 (361-364) | $^{355}$LANNRD***PIVI***SDSPPASPHR$^{374}$ |
| SLS-5 (651-654) | $^{646}$SGVSS***VVAL***SPYVNK$^{660}$ |
| SLS-6 (667-670) | $^{660}$KTITGQC***LPIL***DMETGN$^{676}$ |
| SLS-7 (681-684) | $^{673}$ETGNIGAY***VVLV***DQTGNMATR$^{693}$ |

*SLS-2,3,4 and 5 had a tryptophan added at the N-termini to obtain UV-visibility. SLS1, 7 had an additional glutamic acid added at the N-termini to improve solubility.

**Table S2.** Hydrophobic contacts at the SUMO2/SLS4 interface

| SLS4 residue | SUMO-2 residue | No. of Contacts |
|---|---|---|
| Ile362 | Val30 | 1 |
| Ile362 | Phe32 | 4 |
| Ile364 | Ile34 | 1 |

**Table S3.** Hydrophobic contacts at the SUMO1/SLS4 interface

| SLS4 residue | SUMO-1 residue | No. of contacts |
|---|---|---|
| Ile362 | Ile34 | 1 |
| Ile362 | Phe36 | 1 |
| Ile364 | Phe36 | 2 |
| Ile364 | Leu47 | 2 |

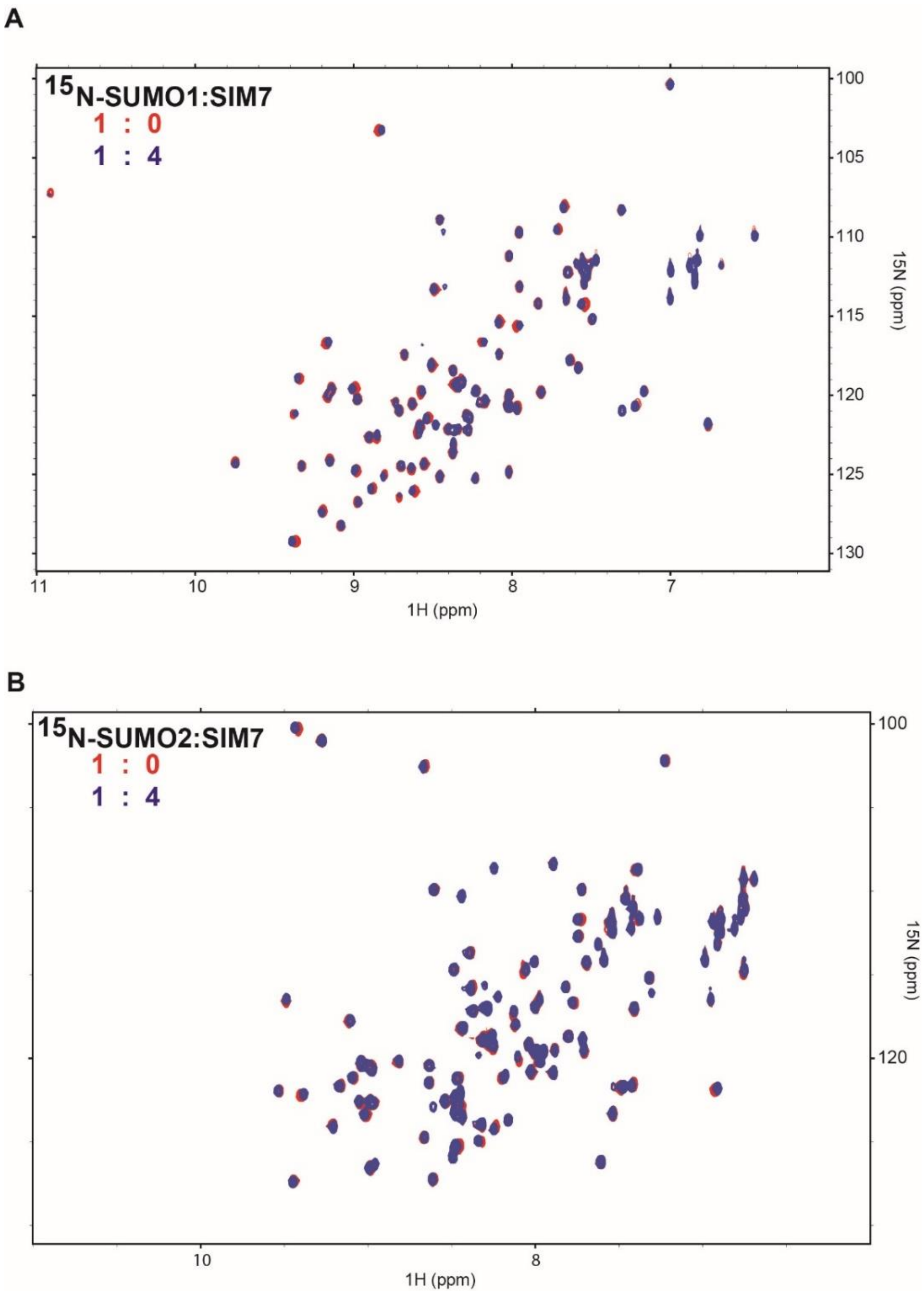


**Figure S1**: (A) Overlay of $^{15}$N-edited HSQC spectra of free SUMO1 (red) and SUMO1 titrated with SIM7 at 1:4 ratio (blue). (B) Overlay of $^{15}$N-edited HSQC spectra of free SUMO2 (red) and SUMO2 titrated with SIM7 at 1:4 ratio (blue).

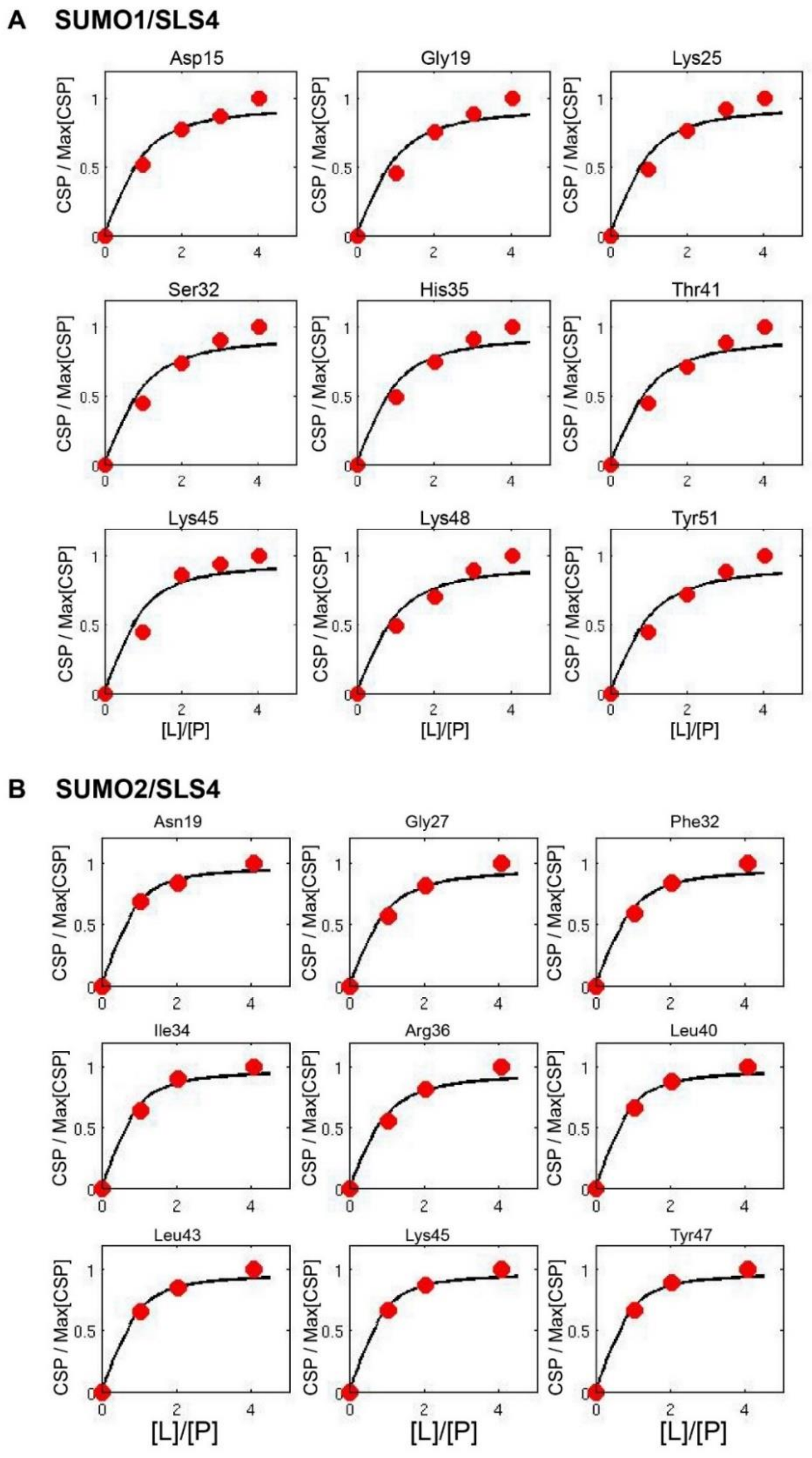


**Figure S2**: (A) Fit of SUMO1 peak shifts against the concentration ratio [SLS4]/[SUMO1] yielded the $K_d$ of the SUMO1/SLS4 complex. (B) The fit of SUMO2 peak shifts against the concentration ratio [SLS4]/[SUMO2] yielded the $K_d$ of the SUMO2/SLS4 complex. The fit of nine typical residues is shown in A and B.

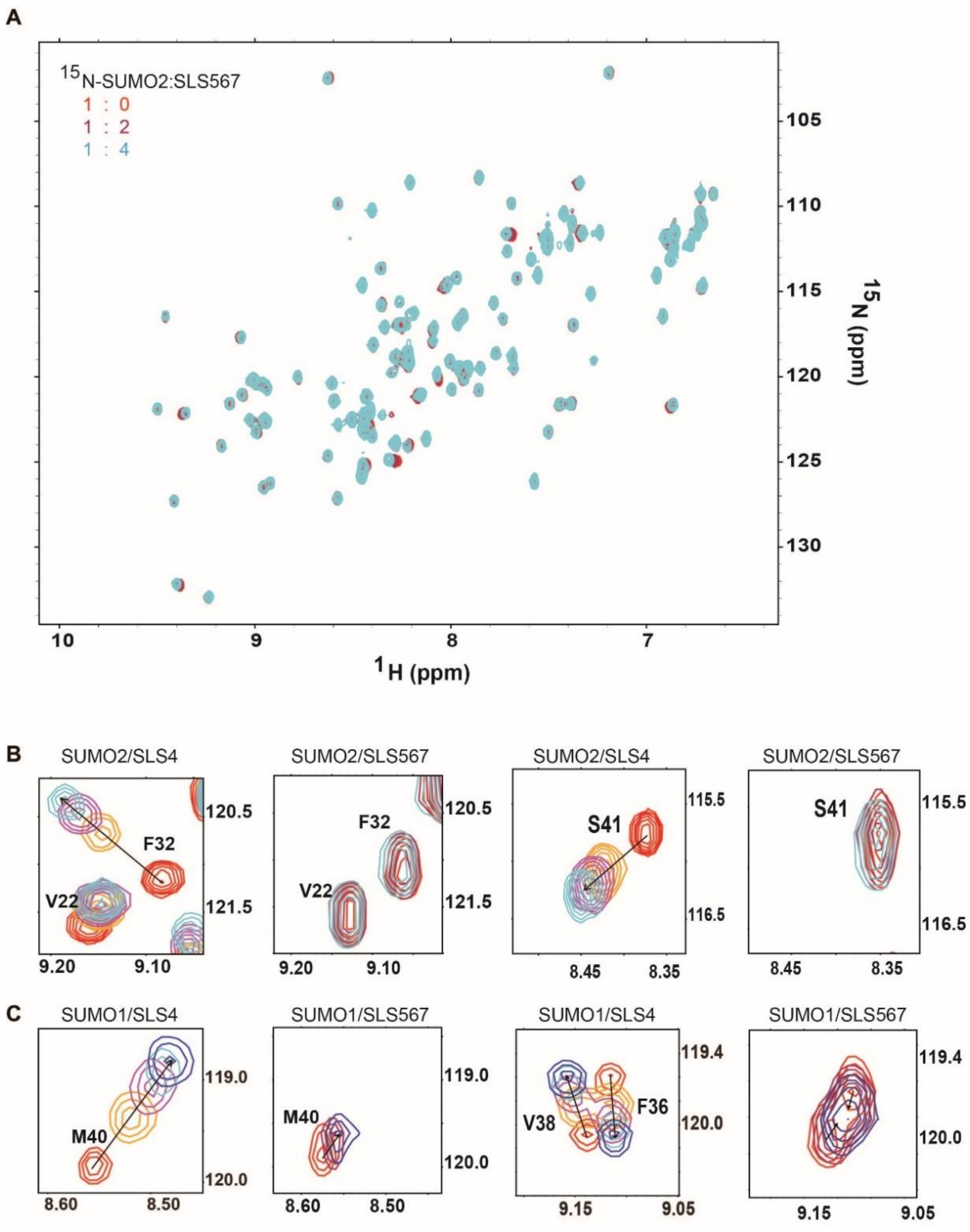


**Figure S3**: (A) Overlay of $^{15}$N-edited HSQC spectra of free SUMO2 (red) and SUMO2 titrated with SIM567 at 1:2 ratio (purple) and 1:4 ratio (cyan). (B) Comparison of peak shifts upon titration for SUMO2/SLS4 and SUMO2/SLS567 for the residues that are present at the SUMO2/SIM interface. (C) A similar comparison as in (B) for SUMO1/SLS4 and SUMO1/SLS567 for the residues that are present at the SUMO1/SIM interface.

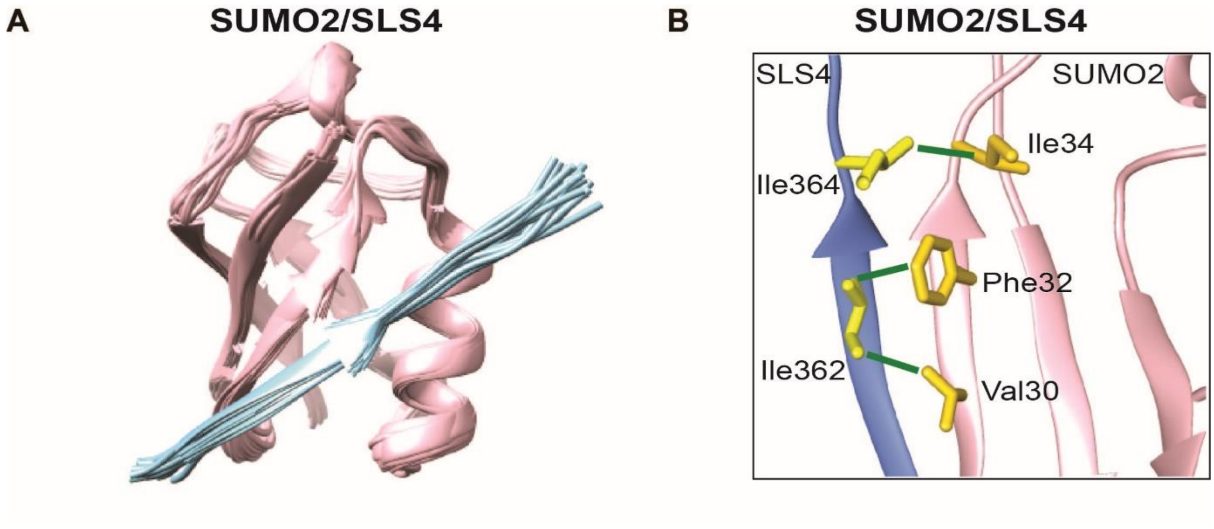


**Figure S4:** (A) Twenty lowest energy solution structures of SUMO2/SLS4. SUMO2 is colored pink, and SLS4 is colored light blue. (B) The hydrophobic contacts between SUMO2 and SLS4 are shown. Side-chains are colored yellow (SLS4) and gold (SUMO2). Green lines mark the hydrophobic contacts.

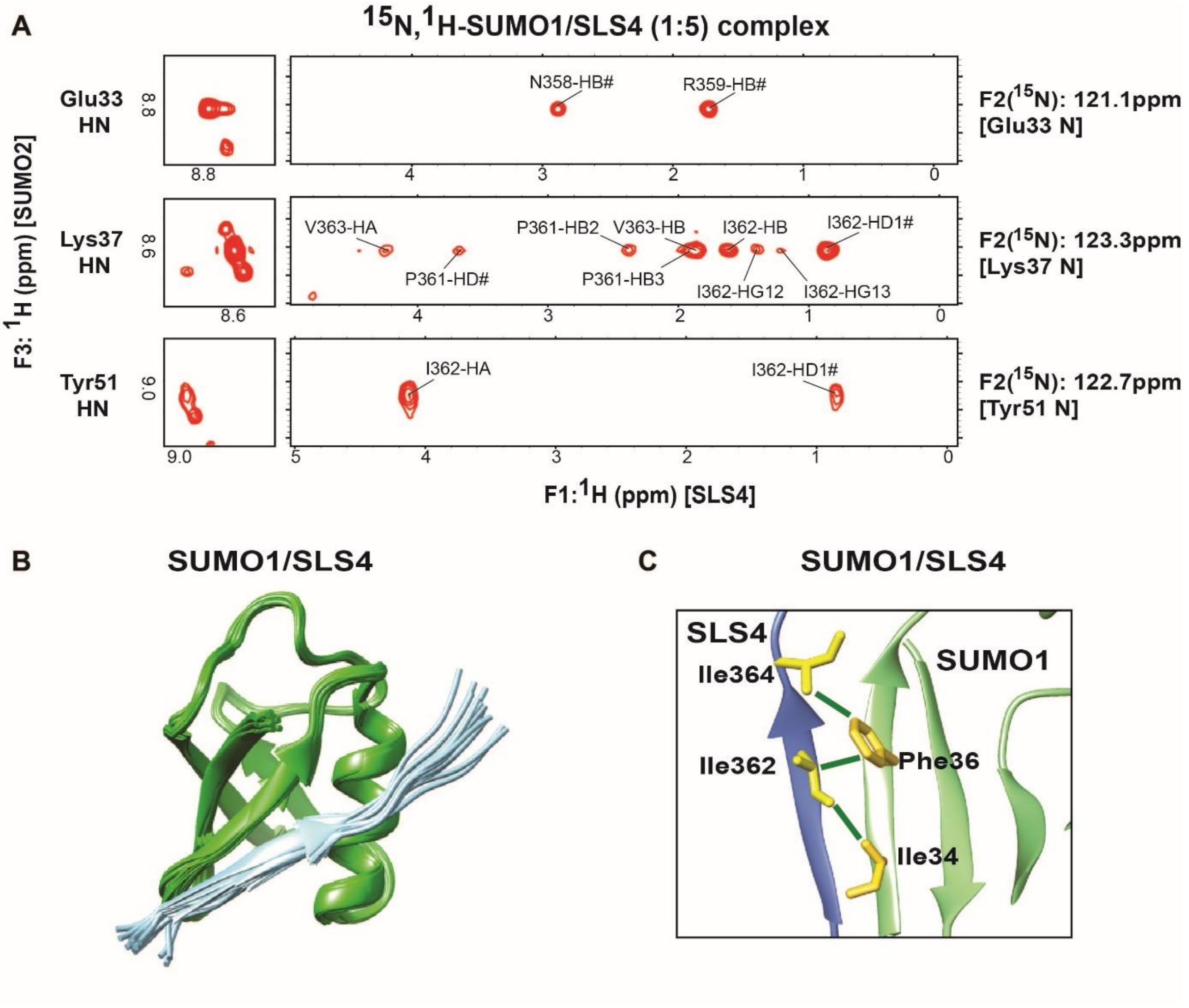


**Figure S5:** (A) The spectral strips from the $^{1}H$-$^{1}H$ plane of $^{15}N$-edited NOESY-HSQC of the $^{15}N$,$^{2}H$-SUMO1/SLS4 (1:5) complex. (B) Twenty lowest energy structures obtained from the structure calculation of SUMO1/SLS4 complex. SUMO1 is colored in green and SLS4 in light blue. (C) Hydrophobic contacts between SUMO1 and SLS4 are marked with green lines. Side-chains are shown in yellow (SLS4) and gold (SUMO1).

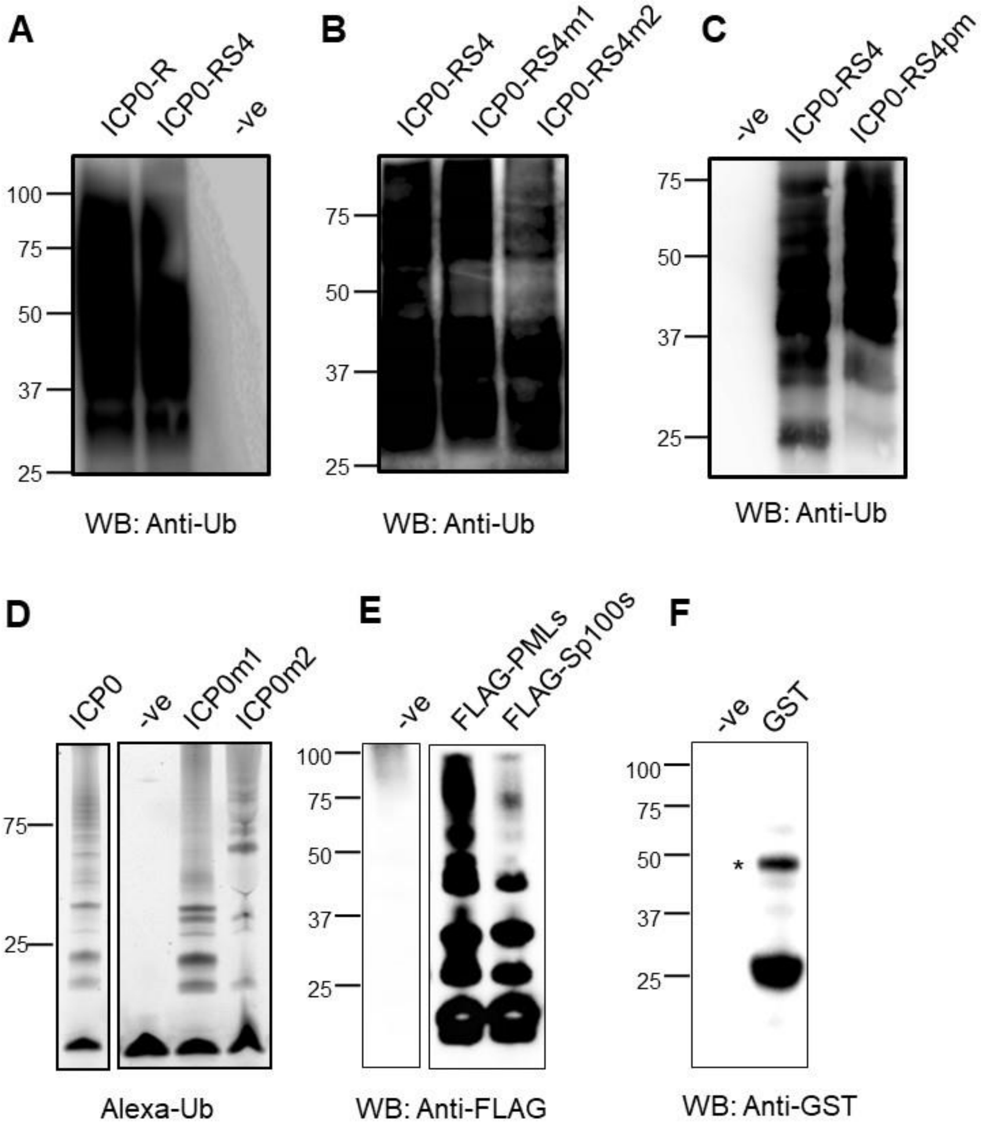


**Figure S6:** *In-vitro* polyubiquitination reaction using ICP0-R, ICP0-RS4, and its mutants as E3s. (A) Both ICP0-R and ICP0-RS4 can assemble polyubiquitin chains with similar efficiency. (B) ICP0-RS4 and its mutants have comparable ubiquitination activity. (C) ICP0-RS4 and phosphomimetic mutant ICP0-RS4m3 exhibit comparable ubiquitination activity. (D) The ubiquitination activity of ICP0, ICP0m1, and ICP0m2. The (–ve) lanes lack ATP. AlexaFluro-Ub was used in this reaction. (E) Post the SUMOylation reaction with FLAG-PMLs and FLAG-SP100s, the SUMOylated proteins are collected in Anti-FLAG agarose beads, washed (3x), separated on SDS gel and blotted with Anti-FLAG antibody. The (-ve) lane does not include a substrate. (F) Post the SUMOylation reaction with GST, the SUMOylated proteins are collected in GSH agarose beads, washed (3x), separated on SDS gel and blotted with Anti-GST antibody. The (-ve) lane does not include a substrate. The band with an asterisk is a dimer of GST.

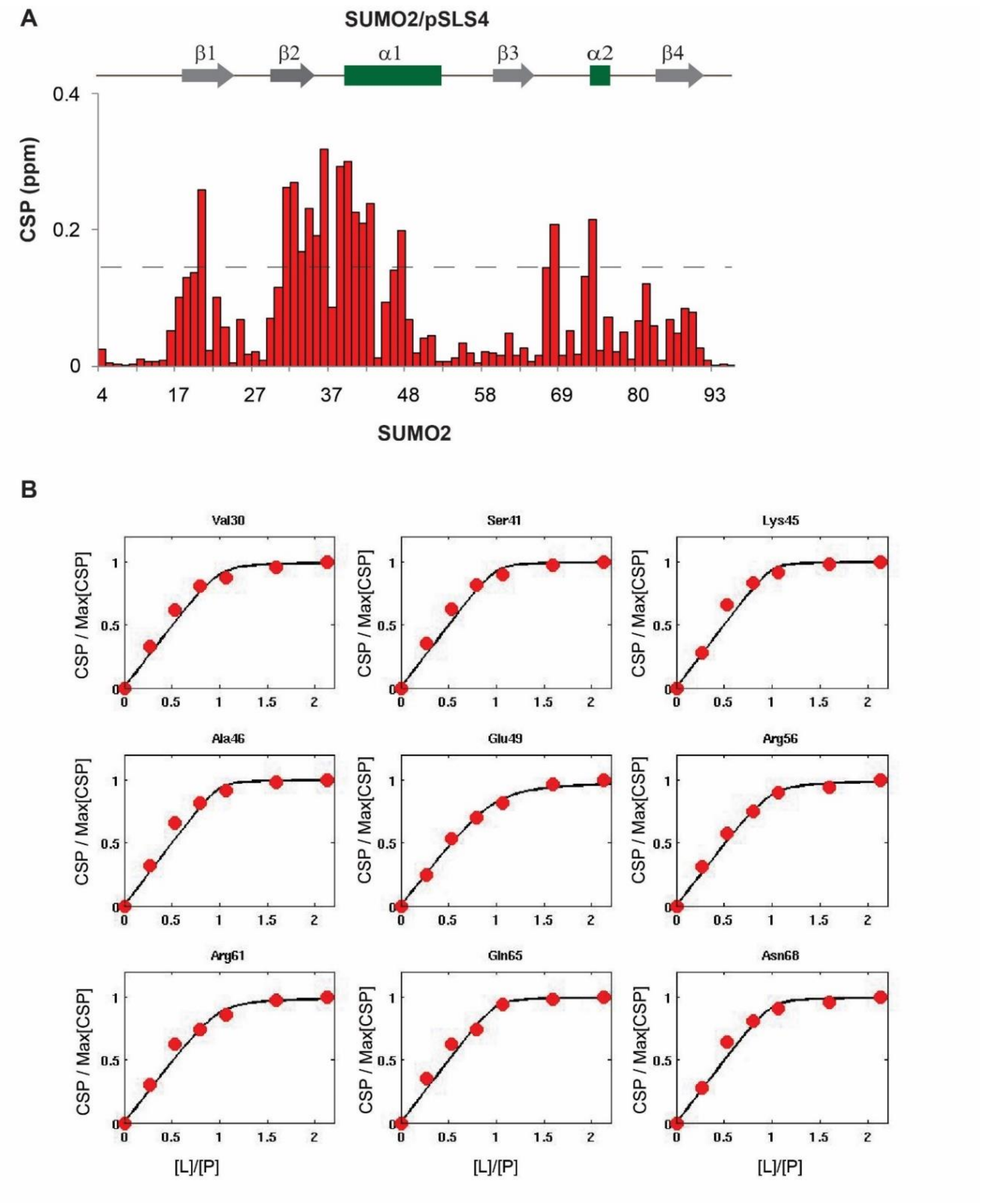


**Figure S7:** (A) The CSP observed in SUMO2 upon titration with pSLS4. (B) The fit of SUMO2 peak shifts against [pSLS4]/[SUMO2] ratio yielded the $K_d$ of the SUMO2/pSLS4 complex. Typical curves and fit of nine residues are shown.

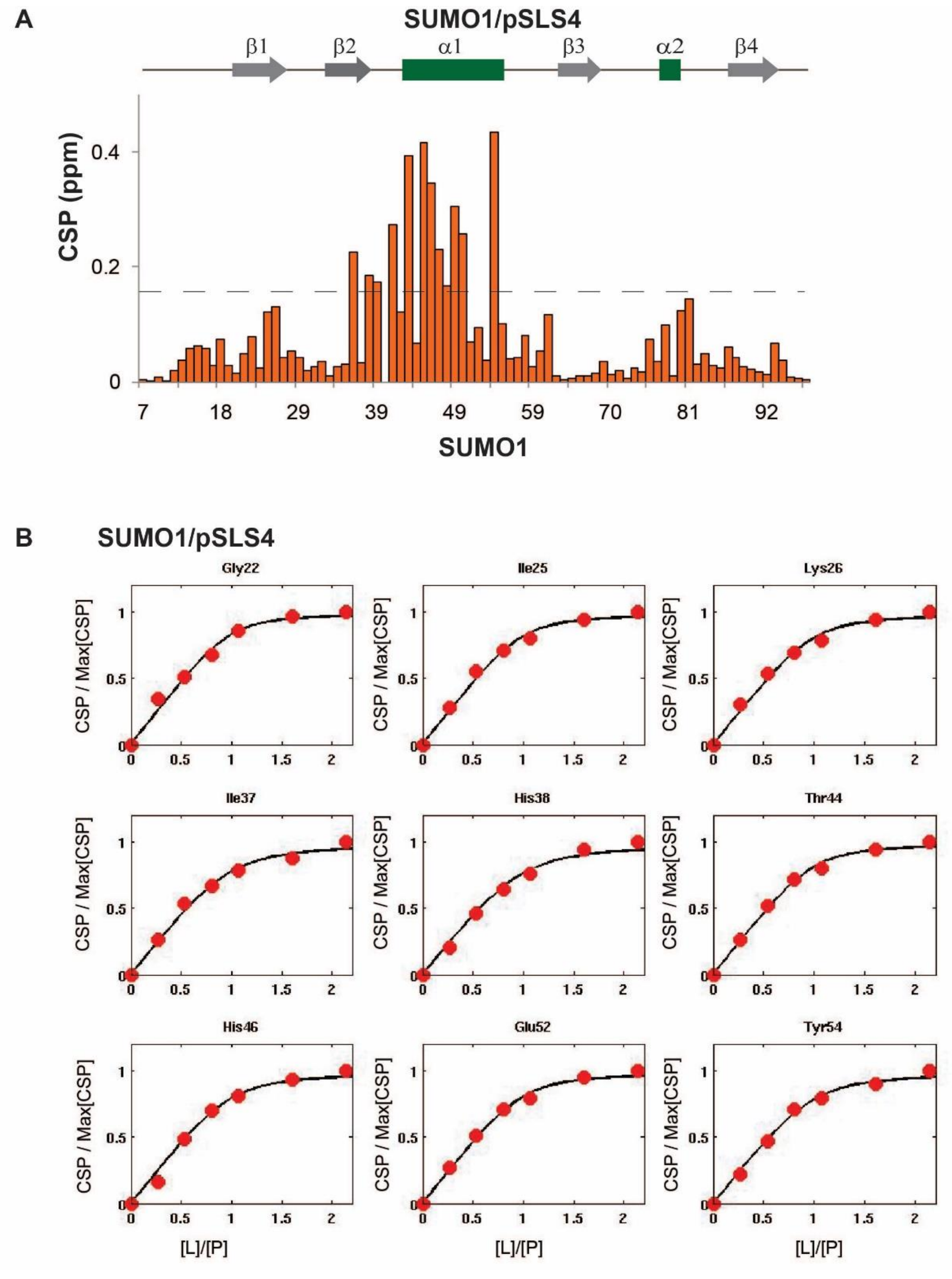


**Figure S8:** (A) The CSP observed in SUMO1 upon titration with pSLS4. (B) The fit of SUMO1 peak shifts against [pSLS4]/[SUMO1] concentration yielded the $K_d$ of the SUMO1/pSLS4 complex.

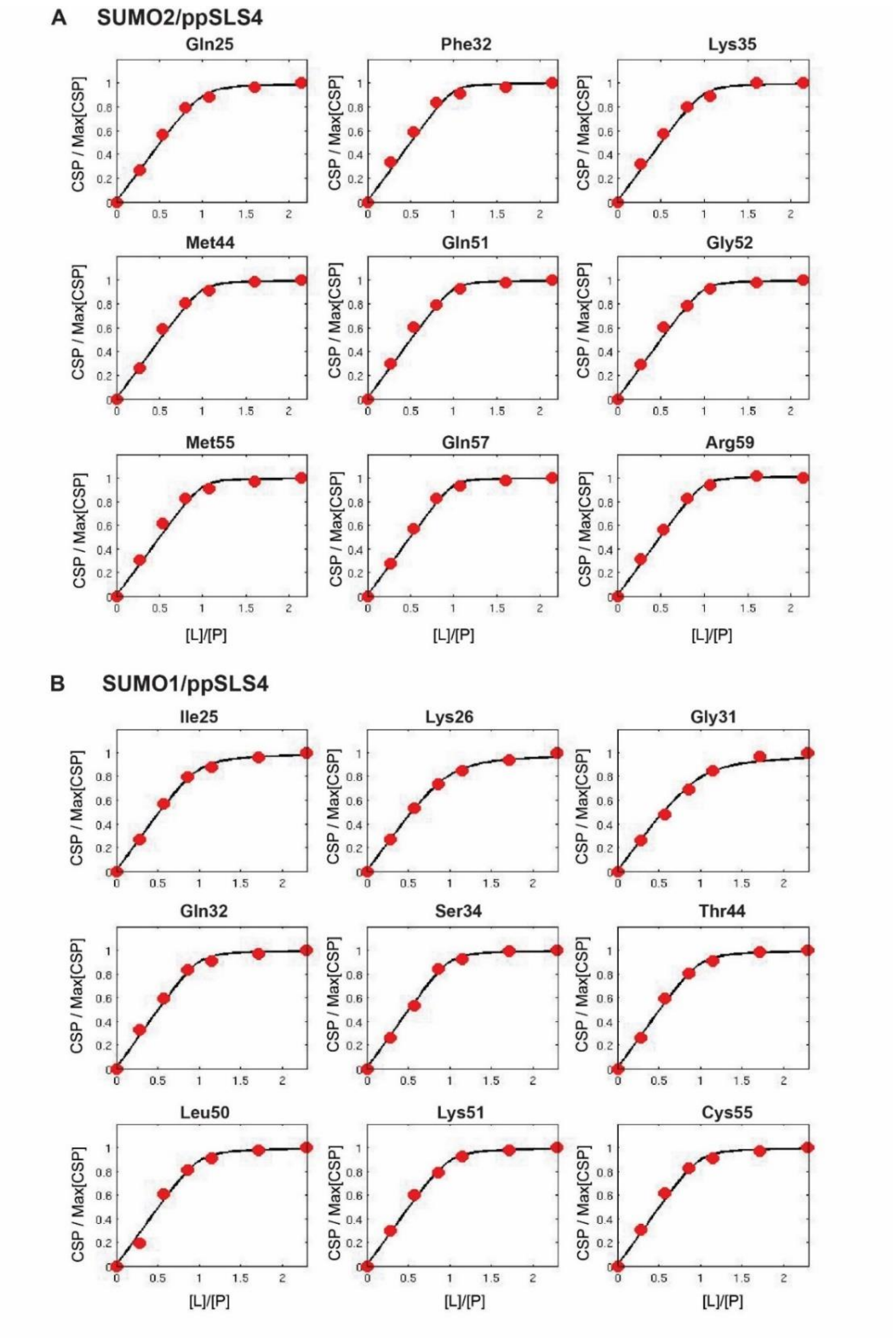


**Figure S9:** Typical curves and fits of SUMO2 and SUMO1 chemical shifts that determined the $K_d$ of (A) the SUMO2/ppSLS4 complex and (B) the SUMO1/ppSLS4 complex, respectively.

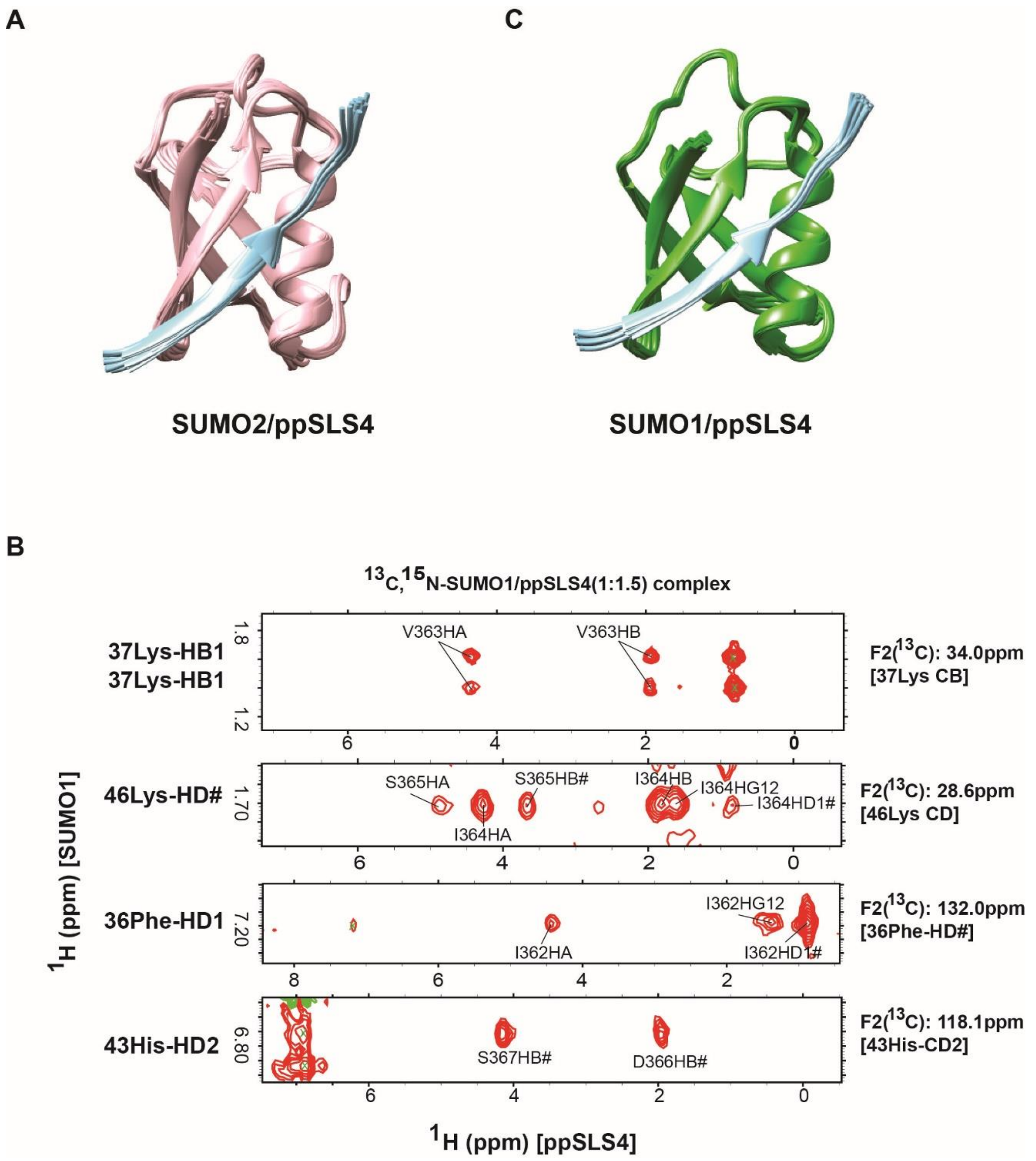


**Figure S10:** (A) Twenty lowest energy structures of the SUMO2/ppSLS4 complex. SUMO2 is colored in pink and SLS4 is colored in light blue. (B) Strips from the $^{1}$H-$^{1}$H plane of $^{13}$C,$^{15}$N-filtered (F1), $^{13}$C,$^{15}$N-edited (F2) NOESY HSQC of the $^{15}$N, $^{13}$C-labeled SUMO1/ppSLS4 (1:1.5) complex. (C) Twenty lowest energy structures from the structure calculation of SUMO1/ppSLS4. SUMO1 is colored in green and SLS4 in light blue.

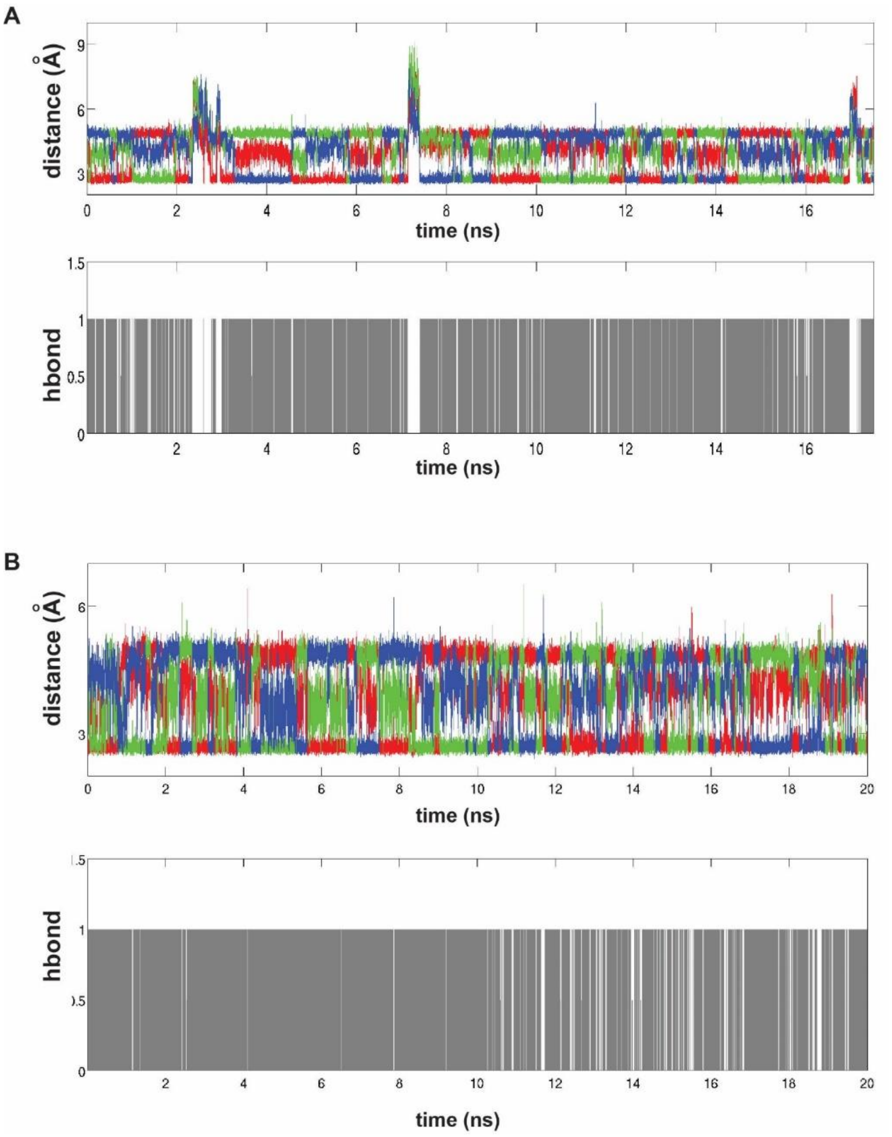


**Figure S11:** Molecular dynamics simulations of the SUMO/SLS4 complexes. (A) Above: Distances between the three phosphorus atoms of phosphorylated pSer367 and Lysine 42 Nζ of SUMO2 are plotted against time. Below: The presence and absence of the hydrogen bond between any one of the phosphorus atoms of pSer367 and Lysine 42 Nζ are digitized to a two-stage process (0 or 1) and plotted against time. (B) Above: Distances between the three phosphorus atoms of phosphorylated pSer367 and Lysine 46 Nζ of SUMO1 are plotted against time. Below: The presence of hydrogen bond between any one of the phosphorus atoms of pSer367 and Lysine 46 Nζ digitized and plotted against time.

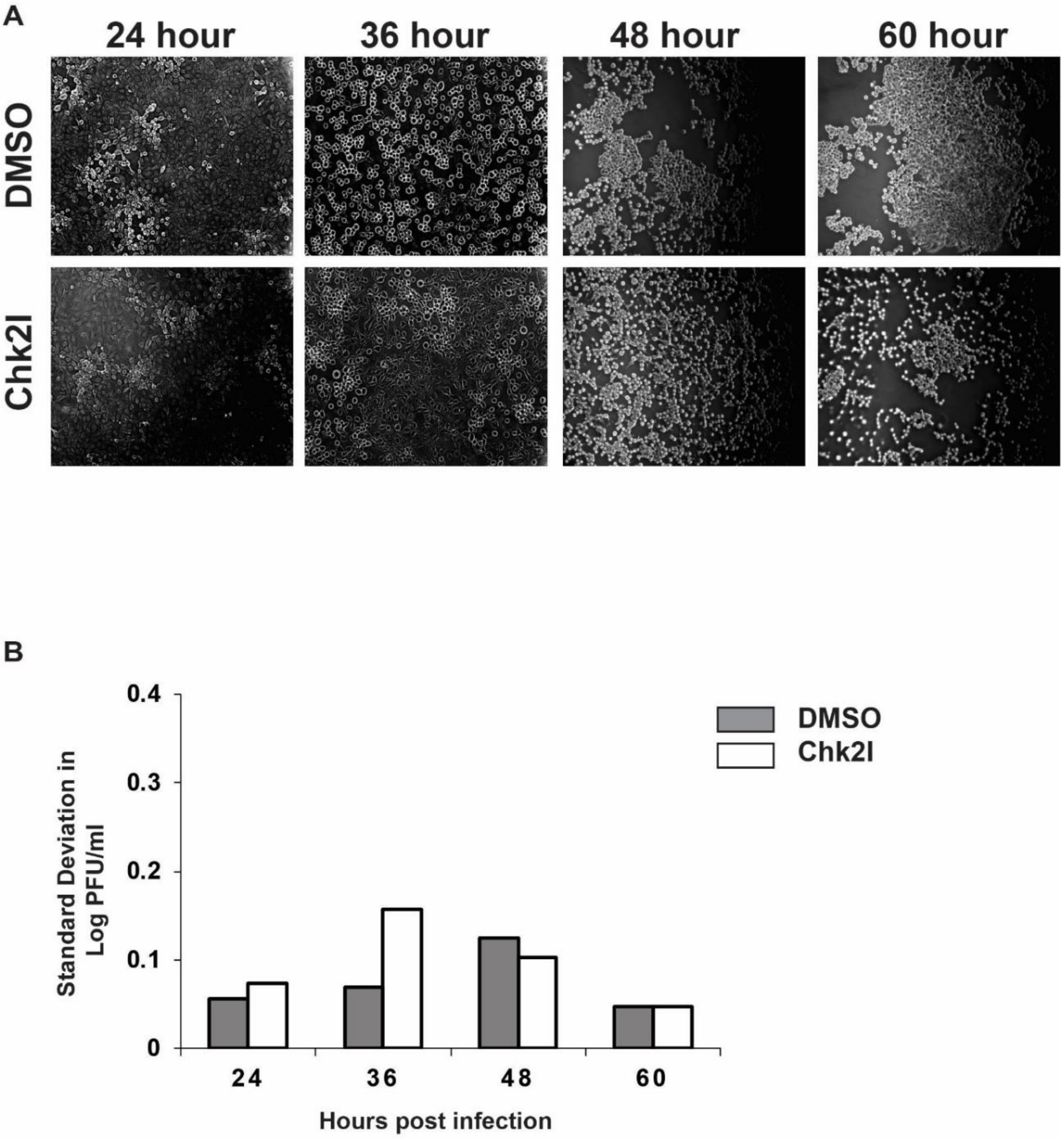


**Figure S12**: (A) HSV-1 infection in the presence of Chk2 inhibitor delayed the progression of (Cytopathic Effect) CPE in the Vero cell line. The Vero cell line was infected at MOI 0.1 in the presence of Chk2 inhibitor and imaged at the indicated time points to monitor the progression of CPE. The untreated control cells (DMSO) exhibit rapid syncytium formation compared to the Chk2 inhibitor-treated cells at 48 hours post-infection. At 60 hours post-infection, the control cells have formed large syncytia while the Chk2I treated cells are still at early stages of syncytium formation. (B) The standard deviation of viral plaque assay data at different hours post-infection.

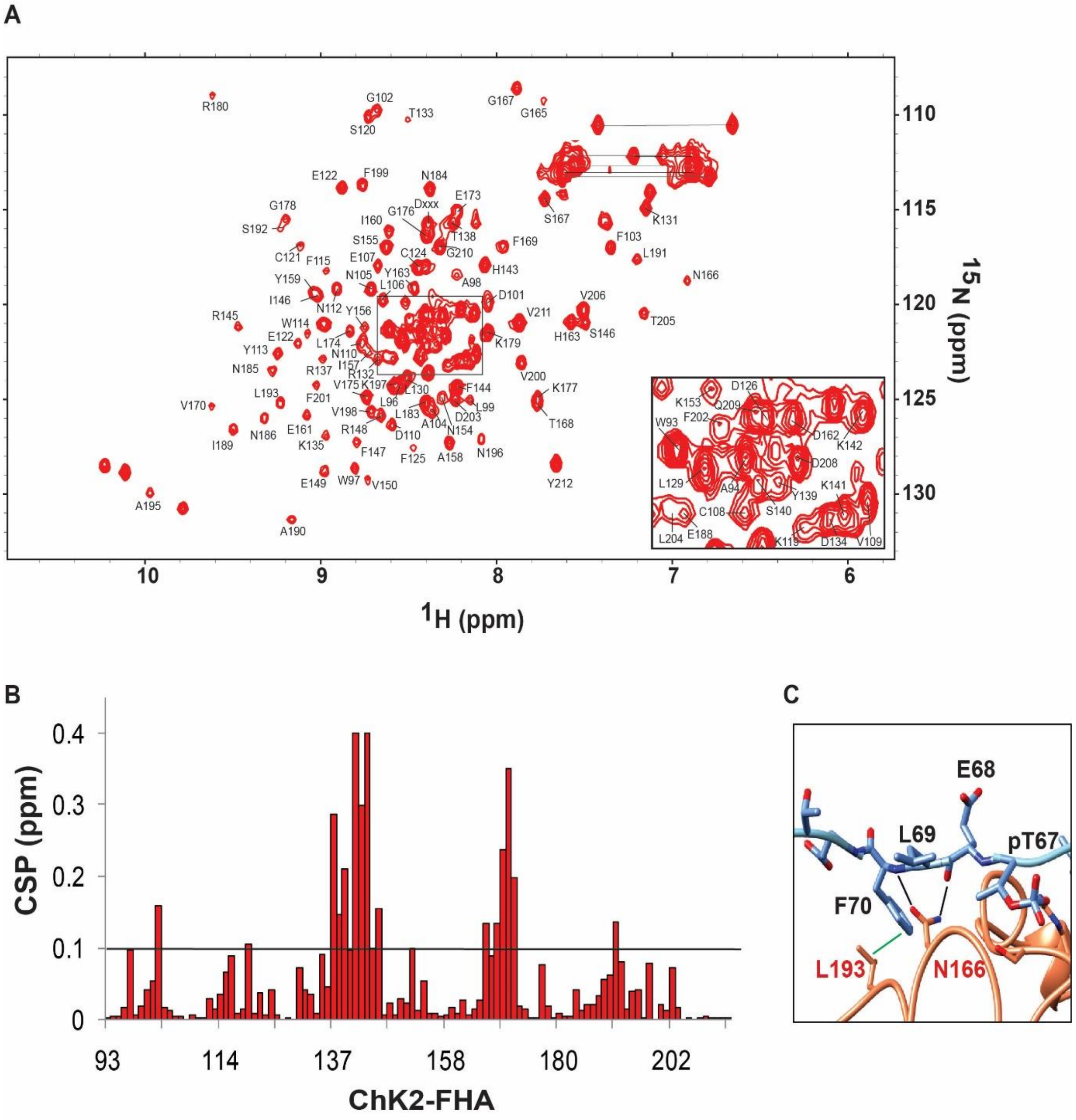


**Figure S13:** (A) Two-dimensional $^{1}$H-$^{15}$N-HSQC spectrum of Chk2-FHA domain. Obtained sequential assignments are indicated by the one-letter amino acid code and residue number. Side-chain amides of glutamine and asparagine are connected by black lines. (B) CSP observed in Chk2-FHA upon titration with the ICP0-FBM. The dashed line corresponds to 2xSD. (C) The contacts observed between Chk2-FHA and ICP0-FBM, apart from the ones involving pT67. Hydrogen bonds are shown in black, and hydrophobic contacts are shown in green.

A

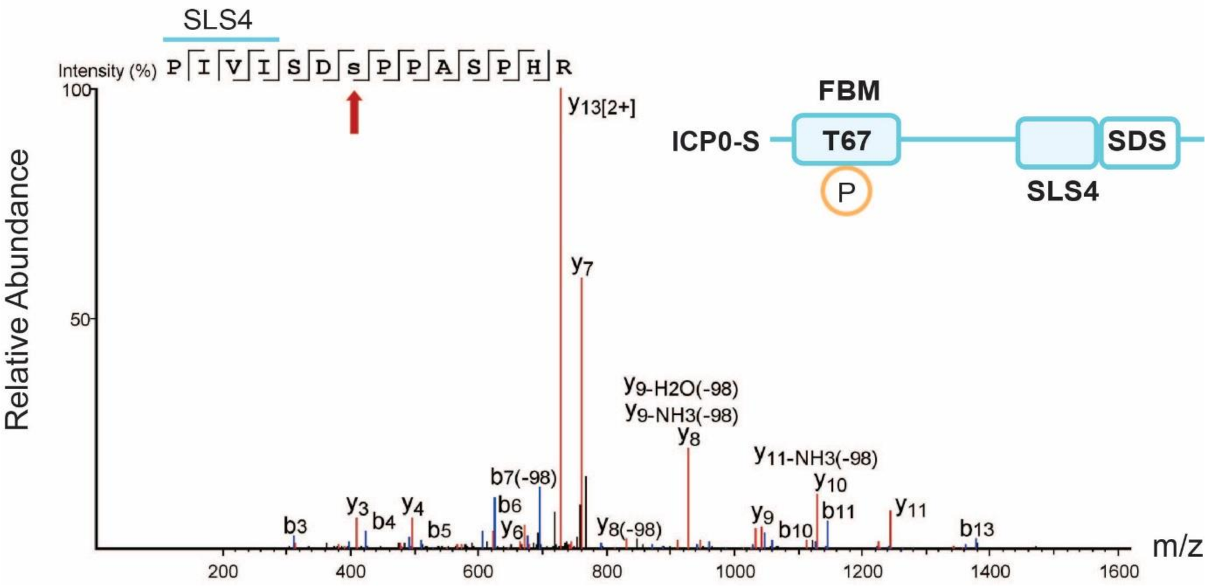


**Figure S14:** MS/MS analysis of the tryptic peptides derived from ICP0-S. Amino acid sequences are indicated with the detected b- and y-type series ions. Ions with neutral loss phosphate (-98 Da) are indicated. MS/MS spectrum corresponded to the tryptic peptide of ICP0-S carrying a phosphorylation moiety at the site Ser367.
